# Antagonism in olfactory receptor neurons and its implications for the perception of odor mixtures

**DOI:** 10.1101/204354

**Authors:** Gautam Reddy, Joseph Zak, Massimo Vergassola, Venkatesh N. Murthy

## Abstract

Natural environments feature mixtures of odorants of diverse quantities, qualities and complexities. Olfactory receptor neurons (ORNs) are the first layer in the sensory pathway and transmit the olfactory signal to higher regions of the brain. Yet, the response of ORNs to mixtures is strongly non-additive, and exhibits antagonistic interactions among odorants. Here, we model the processing of mixtures by mammalian ORNs, focusing on the role of inhibitory mechanisms. Theoretically predicted response curves capture experimentally determined glomerular responses imaged by a calcium indicator expressed in ORNs of live, breathing mice. Antagonism leads to an effective “normalization” of the ensemble glomerular response, which arises from a novel mechanism involving the distinct statistical properties of receptor binding and activation, without any recurrent neuronal circuitry. Normalization allows our encoding model to outperform noninteracting models in odor discrimination tasks, and to explain several psychophysical experiments in humans.

## Introduction

The olfactory system, like other sensory modalities, is entrusted to perform certain basic computational tasks. Of primary importance is the specific identification of odors and the recognition of isolated sources or objects in an olfactory scene. A typical scene in a natural environment is complex: the olfactory landscape is determined by the chemical composition of odorants released by the objects, the stoichiometry of the mixture and the physical location of the objects relative to the observer. An efficient olfactory system is expected to eliminate irrelevant background components and de-mix contextually relevant components received as a blend [1–16].

The importance of filtering a complex background is shared by the olfactory and the adaptive immune systems. In the latter, lymphocytes must quickly and accurately identify a small fraction of foreign ligands in a sea of native ligands [17]. Inhibitory feedback plays a key role in meeting the challenge of a proper combination of rapidity, sensitivity and specificity [18]. The hallmark of inhibitory feedback is a strongly nonlinear response to mixtures of antigens. The advantage of inhibition is the taming of a dominating background; its drawback is the reduction in response amplitudes [18,19].

Antagonism in olfactory receptor neurons has been observed in experiments [20–22], although it has not been quantified systematically. Extensive evidence of intensity suppression and overshadowing in the perception of odorant mixtures comes from psychophysical observations [16,23–25]. The importance of peripheral interactions in shaping mixture perception has been directly shown by electrophysiological and psychophysical measurements [26–28]. However, the functional role, if any, of inhibition at the ORN level remains unknown.

While the above arguments are generally valid qualitatively, their specific application to immunology involves single lymphocytes. This fundamentally precludes a direct analogy to ORNs, which display broad sensitivities and promiscuous binding of a single odorant. The axons of ORNs of a common subtype converge onto glomeruli, where the axon terminals form synaptic contacts with mitral and tufted (M/T) projection neurons leading to the cortex, as well as local periglomerular (PG) interneurons. Discriminatory computations analogous to those performed by individual lymphocytes in the early immune response will be carried out by brain regions such as the olfactory cortex, which receive global information from the ORN ensemble. To achieve a quantitative description of ORN inhibitory effects, it is then imperative to take their global nature into account. In other words, it is necessary to address the knowledge gap between the mixture response properties of a single ORN, the ensemble glomerular response, and ultimately its influence on odor discrimination and perception, which constitutes the goal of the present work.

Previous computational models that examined discrimination tasks have, for simplicity, assumed a linear summation model of mixture response at the ORN level [4,29–33]. Conversely, our emphasis is on explicitly characterizing the ORNs’ biophysical attributes, with a focus on mixture response properties. Our resulting biophysical model is compared to extensive experimental data obtained from imaging glomerular responses to single odorants and their mixtures. A key aspect of the model is that odorant-receptor interactions depend on two distinct features: the sensitivity to binding and the efficiency of activation after binding, respectively. Competitive antagonism occurs when a component in a mixture that binds strongly, activates the downstream transduction pathway less effectively compared to other components. While this might naïvely seem disadvantageous, we show how an antagonistic encoding model has inherent normalization properties, leading to superior performance in odor discrimination and identification tasks. Finally, we make an explicit connection between a variety of psychophysical observations related to the perception of odorant mixtures and inhibitory effects at the single ORN level, providing a neurobiological basis to perceptual phenomena.

## Results

### Biophysics of mammalian olfactory receptor neurons

Odorants in the nasal cavity are captured by G-protein-coupled-receptors located on the cilia of ORNs. The conversion of chemical binding events into transduction currents in the cilia leads to spike signals transmitted to the brain (see Refs. [34,35] for reviews). The signal transduction pathway is complex; here, we build a chemical rate model of the mammalian ORN meant to capture its major response properties. The model complements and extends previous work on mixture interactions [36,37].

In Fig. 1A, we illustrate the transduction pathway as modeled (for details, see Methods). For an odorant *X*, its interactions at the receptor level are represented by a two-step process:

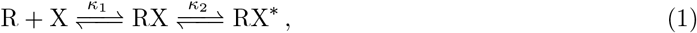

where *κ*_1_ and *κ*_2_ are the ratios of backward to forward rates for the binding and the activation steps. R, RX and RX* represent unbound, bound inactive and bound activated receptors, respectively. Activated complexes lead to the production of cAMP molecules, which diffuse locally and cooperatively open cyclic-nucleotide-gated (CNG) channels permeable to Ca^2^+ and Na^+^ ions. Ca^2+^ ions open Cl^−^ channels further downstream, which produces an outward amplifying Cl^−^ current [39]. Spike firing, in proportion to the current, follows.

**Figure 1:**
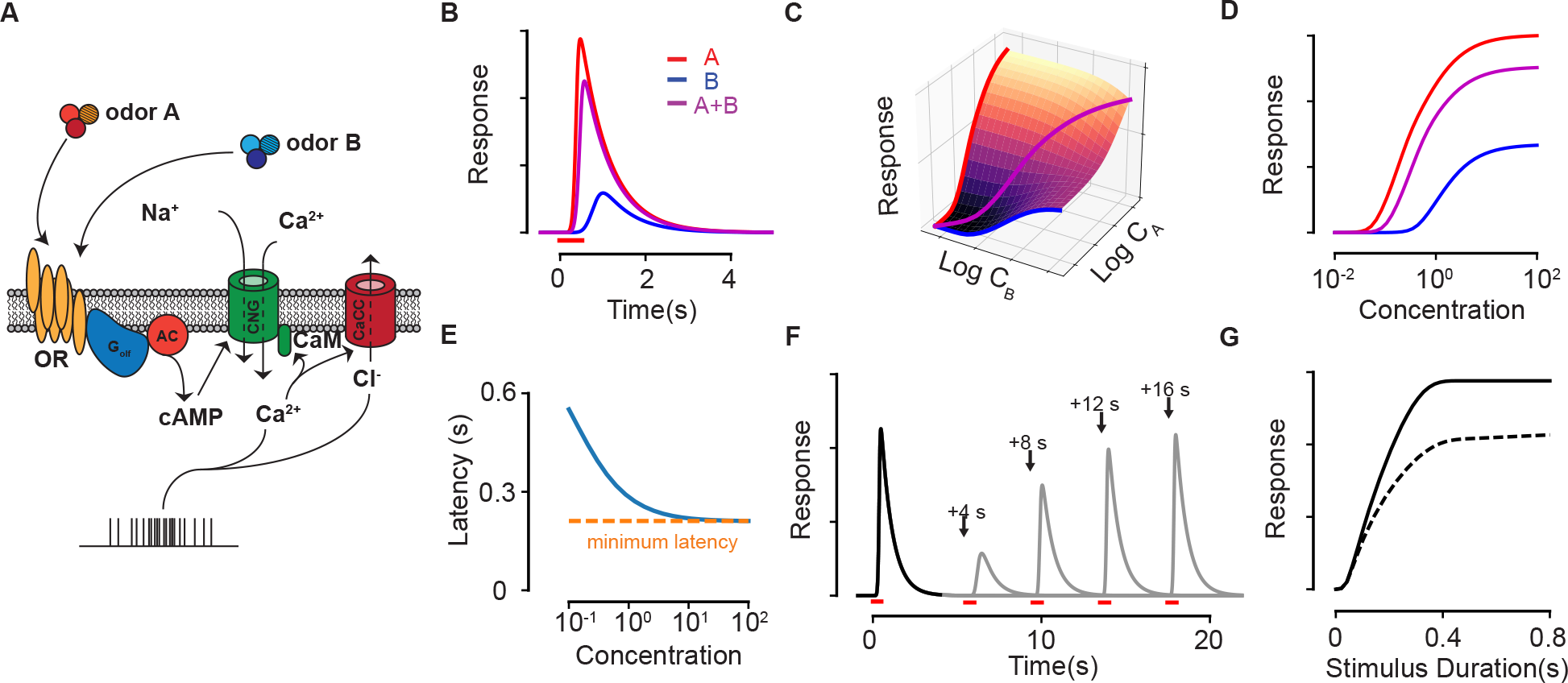
Response properties of an ORN in our biophysical model. (A) Scheme of the modeled ORN signal transduction pathway (adapted from [38]). (B) The temporal responses of an ORN to odorants *A* (red) and *B* (blue) delivered separately and as a mixture (magenta). (C-D) The peak response of the mixture for different concentrations of *A* and *B*. The three colored curves in (C) are plotted separately in (D) on a single axis, whose scale corresponds to *C*_*A*_, *C*_B_ and *C*_*A*_ + *C*_*B*_ for *A*, *B* and the mixture *A* + *B*, respectively. (E) The response latency *vs* odorant concentration. The red, dashed line shows the minimum possible latency due to the limiting receptor activation and cAMP production steps in the signal transduction. (F-G) The Ca^2+^-based adaptation properties of the biophysical model. In (F) the first odorant pulse at *t* = 0 is followed by a second pulse at each of the four shown times in separate trials. Full response is recovered after a few tens of seconds. (G) The peak response of a pulse (solid) and a second pulse (dashed) delivered 10s later against the pulse duration of both pulses.

The specific identity of odorants plays a role only at the level of receptors in our model (except for masking agents, described below). Odorants are characterized by two parameters (mathematically defined by (15) in the Methods): the sensitivity, 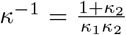, which controls the affinity of *X* to the receptor, and the activation efficacy, *η*, which combines *κ*_2_ with parameters of downstream reaction steps to measure the current produced by *X* once bound. Numerically integrating the set of coupled rate equations presented in the Methods yields the temporal firing rate response of the ORN to pulses of odorant molecules and their mixtures at various concentrations (Fig. 1B-D). For single odorants, the model successfully captures the strongly non-linear peak response for different concentrations of the odorant, the latency in response [40], the quadratic rate of cAMP production [41], and calcium-based adaptation [42] (Figure 1E-G). Since our focus in subsequent analysis will be on the peak response, its form is reproduced here (see Methods for details) for a monomolecular odorant delivered at a concentration *C* for a fixed, short duration:

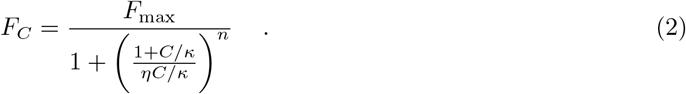

Here, *F*_max_ is the maximum physiologically possible firing rate (which can be rescaled to unity) and *n* is the Hill coefficient. The maximal response at saturating concentrations, *F*∞ = *F*_max_/(1 + *η*^−*n*^) is truncated below *F*_max_ by *η*, which controls the equilibrium level of activated receptors.

On stimulation with more than one chemical species, the different species bind and activate the ORN in distinct ways. Its peak response to a pair of odorant molecules *A* and *B* is a special case of the general formula (14) derived in the Methods, and reads:

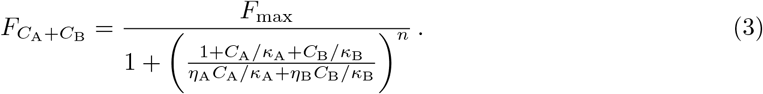

Strong amplification by the ion channels render the mixture response hyper-additive at concentrations close to the sensitivity threshold; at higher concentrations, as the receptors become saturated and the odorants compete for limited binding sites, the response turns hypo-additive. The reduction in response due to competitive antagonism is determined by the binding affinity of the weaker odorant (with lower activation efficacy) relative to the stronger one.

### Experimental validation of the model

The model above captures qualitative and quantitative aspects of receptor neuron dynamics. Before further theoretical analysis, we first present the experimental data used to validate it.

We measured odor-evoked Ca^2+^ signals at ORN axonal terminals within olfactory bulb glomeruli in freely breathing mice. We delivered individual odorants, as well as binary odorant mixtures at different concentrations, spanning four orders of magnitude. For all experiments (see Methods for further details), a craniotomy was performed to gain optical access to both olfactory bulbs in olfactory marker protein (OMP)-GCaMP3 mice [43]. Using a custom-built 2-photon microscope, the field of view typically allowed us to image > 10 glomeruli per field when imaging at 4 Hz (example image in Fig. 2B). After a baseline period, we then delivered a 2 s pulse of single odorants or binary mixtures to the animals through an odorant inlet placed near its nose. At the completion of each experiment, the relative fluorescence ΔF/F signals were then extracted from each glomerulus. For example responses, see Fig. 2B-C.

**Figure 2:**
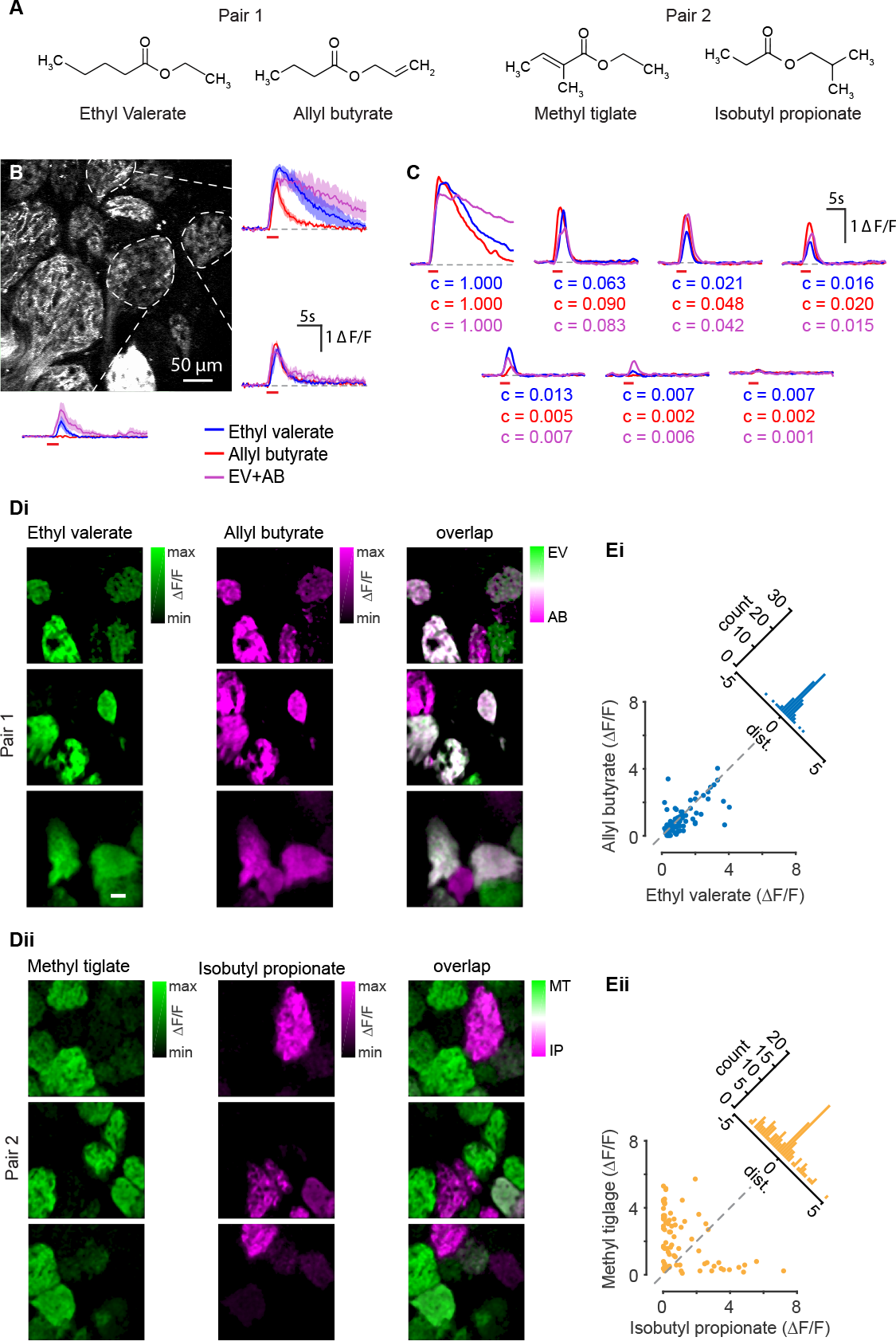
Imaging of overlapping and non-overlapping odorant pairs in ORNs (A) Chemical structure of the four compounds used to elicit Ca^2+^ signals in ORNs. (B) Typical field of view for in vivo 2-photon imaging of ORN odorant responses. The response of three different glomeruli to a 2 s pulse of 10% ethyl valerate (blue line), allyl butyrate (red line) and a binary mixture of both odorants (magenta line). The shaded region shows the variation in response across different trials for the same stimulus. (C) The responses of a fourth glomerulus from the same field as in B across seven concentrations of both ethyl valerate (blue) and allyl butyrate (red) and a mixture of the two (magenta). Concentration values listed below each trace are normalized to the absolute concentration measured at 10% for each odorant and a mixture of both components at 10% each. (Di) Example glomerular responses of three fields for ethyl valerate (left), allyl butyrate (middle), and the degree of overlap between the two (right). Scale bar = 50*μ*m. (Dii) Same as Di but uses a second odorant pair, methyl tiglate and isobutyl propionate. The response pattern for pair one is highly overlapping while the response pattern for odorant pair two remains discrete. (Ei) Scatter plot of the peak response for odorant pair one and the distance from unity (inset) for each glomerulus that responded above threshold (threshold = 0.2530 ΔF/F) at 10% odorant concentration. (Eii) Same as Ei for odorant pair two.

For all experiments one of two odorant pairs were used, here called pair one (ethyl valerate and allyl butyrate) and pair two (methyl tiglate and isobutyl propionate). The odorant pairs were selected based on previous studies demonstrating the similarity or dissimilarity in the spatial pattern of activated glomeruli generated by each of the odorants (see [13], Figs. 2D-E) so that our observations could be extended across diverse blends.

From nine mice, we extracted Ca^2+^ signals from 296 total glomeruli, of which, we included 138 glomeruli for odorant pair one and 108 glomeruli for odorant pair two in our analysis. To quantify the degree of overlap between each odorant in a pair, we calculated the Euclidean distance from unity for the peak odor-evoked response at each glomerulus for both odorants at their highest concentration (10%; Fig. 2E) for all glomeruli responding above a threshold determined by receiver operating characteristic analysis (threshold = 0.25). A distance of zero indicates an equal response to both odors in a pair. The mean distance for pair one was 0.35±0.05 and 1.48±0.12 for pair two (*p* < 0.0001, rank-sum test, *n* = 77 glomeruli for pair 1 and *n* = 89 glomeruli for pair 2). The increased distance measured in pair two indicates that glomeruli were, on average, more selective for each of its constitutive odorants compared with pair one.

Having established two pairs of odorants that resulted in disparate levels glomerular overlap, we then used the response of each glomerulus at discrete concentrations to construct response curves. For our analysis, we calculated the mean of the peak ΔF/F responses of at least three repeats at each odorant and mixture concentration. The parameters *κ* and *η* for each odorant were determined by simultaneously fitting the mean peak response from the individual odorants and the mixture using equations (3) and (2). Response curves were obtained by using the best fit parameters in equation (2) for each odorant (Fig. 3B, solid red and blue lines in Bi and solid orange and teal lines in Bii) and in equation (3) for the mixture (Fig. 3Bi-ii, solid magenta or purple line). The dashed magenta or purple lines in Fig. 3B are direct sigmoidal fit to only the mixture data. The distribution of the parameters *κ* and *η* across all glomeruli are shown in Fig. 3C-D. We verify the broad distribution of *κ*^−1^, which is a central characteristic of olfactory coding [47]. We further observe a broad distribution of *η* values, which implies a range of activation efficacies. The relationship between *κ*^−1^ and *η* for a single ORN, namely their joint statistics, plays a major role in determining the encoding capacity of the ORN ensemble, as detailed in later Sections.

**Figure 3:**
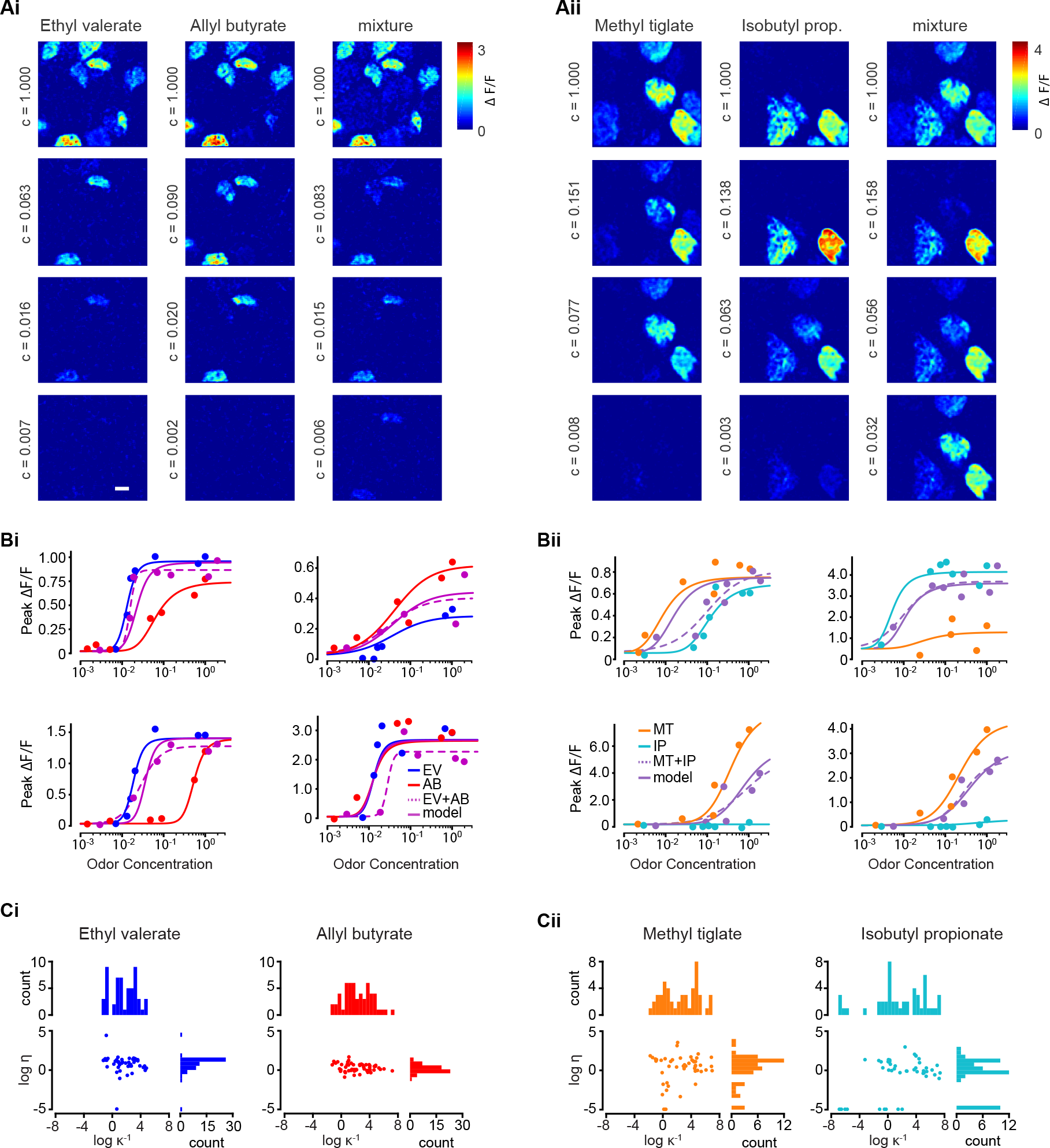
Experimentally obtained ORN-odorant response curves. (Ai-ii) Whole-field ΔF/F responses across four odorant concentrations for odorant pair one and odorant pair two. Scale bar = 50*μ*m (Bi-ii) Example response curves obtained from four individual glomeruli using each odorant pair. Red and blue circles are the mean peak responses for pure odorant compounds, magenta circles are odorant mixtures. The solid red, blue and magenta lines are obtained using the best fit parameters (*κ* and *η*) in equation (2) for each odorant and in equation (3) for the mixture. The dashed magenta line is obtained from a sigmoidal fit to only the mixture data points. In Bii the orange circles and curves are responses to methyl tiglate, teal circles and curves are isobutyl propionate, and purple is the mixture. (Ci-ii) The best fit parameters *κ* and *η* obtained for each glomerulus, and their distributions, for the four odorants used in this study.

Figs. 3Bi-ii demonstrate that our model is indeed capable of capturing qualitative aspects of ORN odorant mixture responses. We make three specific observations: (1) The saturation level is odorant-dependent for the same ORN type as seen in the top two panels of Fig. 3Bi and top right panel of Fig. 3Bii. (2) The mixture response is mostly hypo-additive and is usually inhibitory at higher concentrations, i.e., the mixture response is lower than the most responsive odorant in the pair. This is strikingly seen in the bottom panels of Fig. 3Bii. (3) We observe several instances of non-competitive suppression, such as in the bottom right panel of Fig. 3Bi, where the mixture response is lower than the response of both odorants.

#### Masking

While a purely competitive model of mixture interactions captures most cases, a significant fraction show discrepancies (as noted above), which were also observed in previous experiments with single ORNs [36]. Non-competitive interactions are particularly manifest in synergy or suppression, which correspond to the mixture response curve lying above or below the individual response curves for each odorant, respectively, neither of which is possible with pure competition. Here, we show how non-competitive antagonistic effects, namely masking, can generate those effects. Non-competitive inhibition due to PI3K-dependent antagonism [45] is beyond the scope of this paper.

Masking is the phenomenon of non-specific suppression of CNG channel currents [22,41]. Experimental evidence suggests that masking agents disrupt the lipid bilayer on the cell membrane, and thereby alter the binding affinity of cAMP to the CNG channels [21]. Masking agents can also be odorants (like amyl acetate), i.e. they also bind to receptors and excite the transduction pathway. We suppose that the agents bind to sites on the lipid bilayer, and that multiple masking agents compete for the available sites. Similar to (1), the effects of a masking agent are determined by its affinity *K*_*M*_ for the masking binding sites, and a masking coefficient *μ*, which measures its inhibitory effects once bound (see Methods). The latter quantifies the lowered affinity of cAMP for the CNG channels due to masking by reducing the activation efficacy *η* that appears in (2). It follows that the maximal firing rate is thereby reduced (see (19) in Methods).

The model above (and detailed in the Methods) reproduces qualitative features of odorant suppression observed in experiments [22] (Fig. 4A) and shows a good fit with available experimental data [46] on masking agents (Fig. 4B). Importantly, both suppression and, counterintuitively, synergy are possible mixture interactions that could arise due to masking (Fig. 4C-D). Synergy is qualitatively due to the taming of suppressive effects. For instance, let us consider a component *A* that binds masking sites more weakly than *B*, yet it is a stronger suppressor. If *A* binds and activates the ORN more strongly than *B*, the net effect of mixing *A* and *B* is to reduce the suppressive masking effect of *A* and unmask its strong activation properties, which can exceed the individual response curves as in Fig. 4C.

**Figure 4:**
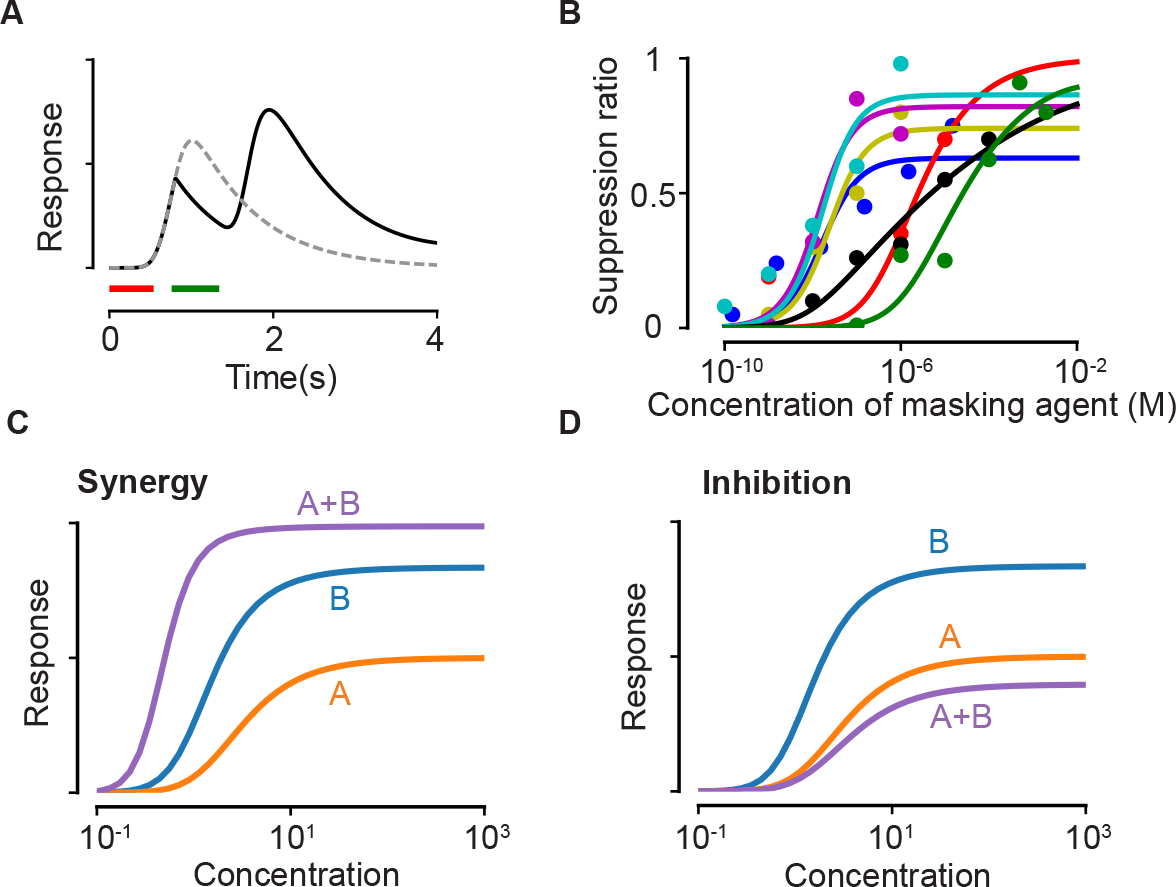
Suppression due to masking. (A) The dashed line shows the response predicted by our model with only the first odorant (delivered during the red time window), while the solid line shows the predicted response when a second pulse of a highly masking odorant is delivered shortly after during the green window (plotted as in [22]). (B) Experimental data on the masking effect of various masking agents (circles) and fits to theory (lines). Blue: 2,4,6-trichloroanisole, red: 2,4,6-tribromoanisole, yellow: phenol, Magenta: 2,4,6-trichlorophenol, cyan: trichlorophenetole, black: L-cis diltiazem, green: geraniol. (C-D) Mixture response curves displaying synergy (C) and inhibition (D). The curves are plotted as in Fig. 1D.

### Olfactory encoding and antagonism

Equipped with the model above, we now proceed to investigate the functional consequences of antagonism. To this end, we first define a model of olfactory encoding that focuses on competitive antagonism and introduce simplifying assumptions to highlight the main ideas.

An odorant is defined by two N-dimensional vectors of sensitivities ***κ*^−1^** and activation efficacies ***η*** across *N* distinct ORN receptor types. We take *N* = 250, which is large enough to generalize our results across species. Parameters for different odorants are drawn independently from log-normal probability distributions (see Methods). The width of the *κ*^−1^ distributions reflects the broad sensitivities of odorants, spanning about six orders of magnitude [47].

An odorant caught in a sniff elicits a glomerular pattern of response whose individual activations vary in magnitude and progress differently in time. We focus on the vector ***y***, which represents the peak responses (2) to the odorant for the different ORN types. The vector of continuous values ***y*** is converted into a binary vector ***z*** by applying a threshold τ, and partitions the glomeruli into two subsets, active and inactive glomeruli (see Fig. 5a). In general, any continuous read-out is demarcated into a few discrete, distinguishable states depending on the level of intrinsic noise in the system ; the threshold allows us to ignore the noise in signal transduction, thus making the analysis simpler.

To characterize the statistical properties of the glomerular binary response ***z***, we introduce two quantities: (i) the sparsity *p*, which is the fraction of glomeruli activated at saturating concentrations of an odorant for a given threshold τ ; (ii) the antagonistic factor *ρ*, which measures the strength of competitive antagonism. To define *ρ*, we note that an odorant *A* competitively antagonizes odorant *B* (at equal concentrations) when its sensitivity exceeds that of 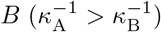, yet its activation efficacy is lower than *B* (*η*_*A*_ < *η*_*B*_). A quantification of this relationship is the Pearson correlation coefficient between binding and activation strengths across the ORN ensemble :

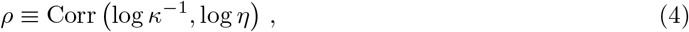

where the logarithms conveniently account for the broad range of the two variables. From (3), we can write the mixture response of an ORN in terms of effective mixture parameters 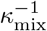 and *η*_mix_:

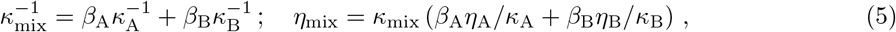

where *β*_*A*_ and *β*_*B*_ are the respective fractions of *A* and *B* in the mixture.

Since sensitivities are broadly distributed, for an equiproportionate mixture we typically have 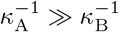 or vice-versa. If we suppose the former :

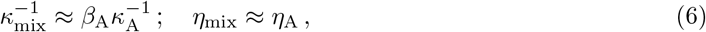

for typical values of *η*_*A*_/*η*_*B*_. Relations (6) generalize to complex mixtures as the width of the sensitivity distribution ensures the dominance of one of the *κ*’S even for relatively large numbers of components.

The key consequence of (6) is that when *κ*^−1^ and *η* are independent i.e., *ρ* = 0, the distribution of *η*_mix_ matches the distribution of *η*_*A*_. It follows that the statistics of activation is conserved as the complexity of the mixture increases, i.e., the population response is “normalized” (Fig. 5C,D). We conclude that normalization of the mixture response, sparse representations and pattern decorrelation, are direct consequences of antagonism in the receptor encoding, independent of any neural circuitry.

**Figure 5:**
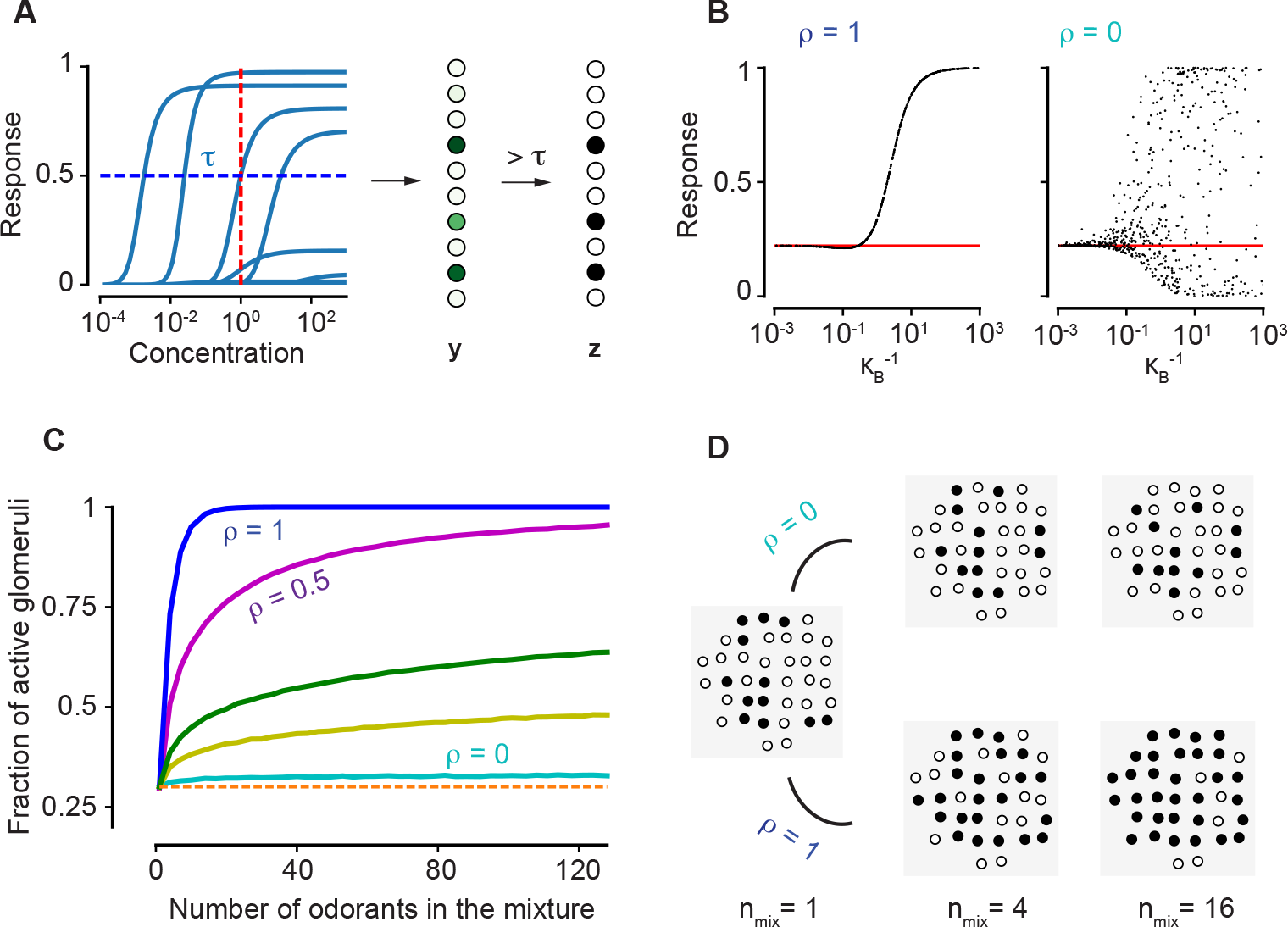
The antagonistic encoding model and normalization. (A) The response curves of a collection of ten ORN types to an odorant. The vector ***y*** of continuous levels of response at a particular concentration (red, dashed line) yields a binary vector ***z*** of activation by imposing a threshold τ (blue, dashed). (B) Our prediction (16) for the response to a mixture of two odorants. The red line indicates the level of the response to odorant *A* alone, which has *κ*_*A*_ = *η*_*A*_ = 1. The concentration of both odorants *A* and *B* is unity. Black points correspond to a sampling of *η*_*B*_ for the value of 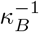 reported on the horizontal axis. A unique value of *η*_*B*_ is obtained when the antagonistic factor *ρ* = 1, as defined by (4). (C) The fraction of active glomeruli as the number *n*_*mix*_ of odorants in the mixture is increased, each of which individually activates 30% of the glomeruli (p = 0,0.1, 0.2,0.5,1 from the bottom to the top curves). (D) The glomerular pattern of activation for *ρ* = 0 and *ρ* = 1 as *n*_*mix*_ increases, shown to contrast the sparsity of their glomerular responses.

The *ρ* = 0 condition of decorrelation between *κ*^−1^ and *η* (sensitivity and activation efficacy, respectively) is important for the argument above since, even though (6) holds generally, the constraint 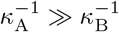 implicit in (6) would otherwise bias the statistics of *η*_*mix*_, which would not coincide with the distribution of individual *η*’s. The extreme case is *ρ* =1, when there is no antagonism as the odorant that binds best also has the strongest activation. This corresponds to an ORN behaving as a logical OR gate, a feature shared by any additive model of mixture response. Structural constraints at the receptor level are likely to hamper a perfect decorrelation *ρ* = 0. However, normalization does not require an exact equality and its effects fade gradually as *ρ* increases (see Fig. 5C). The extent of the advantageous effects of normalization (and the *ρ* value) depends on the sparsity of activation for single odorants, the number of components in the mixture and their properties. Indeed, the effect of normalization on the detection of an odorant in the presence of a large number of other odorants, depends on the balance between two opposing factors. On the one hand, normalization induces an advantageous effect of maintaining sparsity and preventing saturation of the bulb, leading to easier segmentation. On the other hand, the number of glomeruli corresponding to each odorant is greatly reduced, which makes detection harder.

### Performance in discrimination and identification tasks

To explore how our model performs in discrimination tasks, we next compute the performance of the antagonistic encoding model described above in detecting a known odorant from a large background of unknown odorants, i.e., figure-ground segregation [13]. The capacity of an optimal Bayesian decoder in the task depends on the mutual information *I*(*T*; ***z***) (in bits) that the glomerular pattern ***z*** preserves about the presence (*T* = 1) or absence (*T* = 0) of the target. Figure 6A demonstrates that an encoding model with significant antagonism (*ρ* = 0) contains more information than a non-antagonistic model (*ρ* = 1) as the background increases in complexity. For a specified sparsity *p*, we compute the level of antagonism that maximizes information transmission when the background varies both in composition and complexity, such as those experienced in natural environments (see Methods). We find that for experimentally observed levels (0.1-0.3) of sparsity [47–50], it is always advantageous to incorporate non-zero levels of antagonism into odorant encoding (Fig. 6B).

We measure the performance of a linear classifier in component separation, the task of identifying several known components from a mixture, for different levels of antagonism. Component separation is qualitatively different as the information about the other known odorants can be recurrently exploited to extract more information about an odorant’s presence or absence [33]. First, a linear classifier is trained to individually identify 500 known odorants in the presence of other odorants from the set. In the test phase, a mixture which contains 1 to 20 known components, uniformly chosen, is delivered. The hit rate measures the fraction of odorants that were correctly identified, while the false positives (FPs) is the number of odorants out of the 500 that were not actually present but were declared to be present. Generalized Receiver Operating Characteristics (ROC) curves are drawn by varying the detection threshold of the linear classifier for each odorant (Figs. 6C-D). We find again that antagonism in receptors yields superior performance, independent of sparsity.

**Figure 6:**
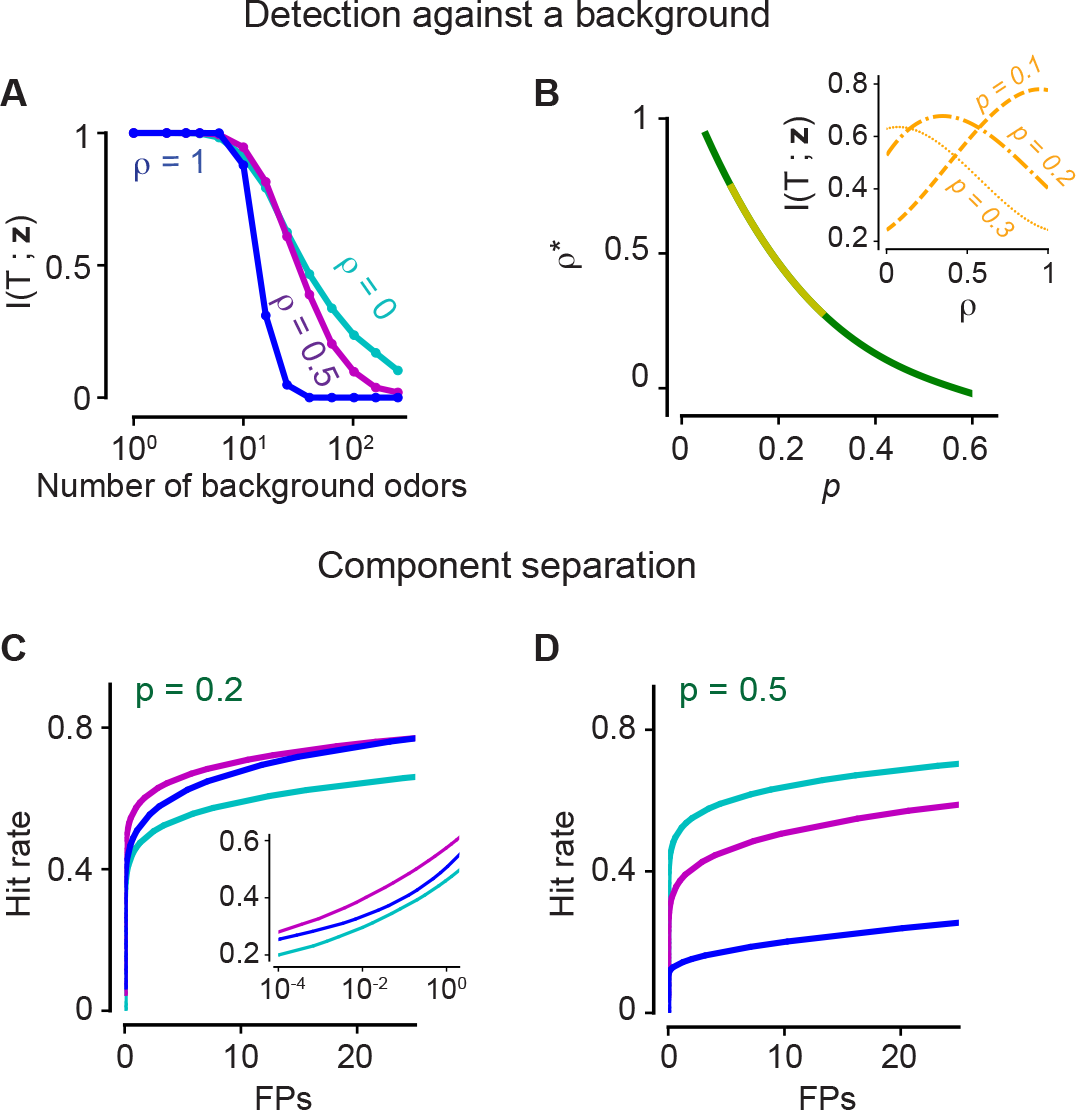
The positive effects of antagonism on odor discrimination and component separation. (A-B) Figure-ground segregation: (A) The mutual information between the presence of a target odorant and the glomerular activation vector ***z*** for the antagonistic factor (14) *ρ* = 0,0.5,1 with varying number of background odorants in the mixture. (B) The optimal value *ρ*\* of the antagonistic factor *ρ* that maximizes mutual information when the number of background odorants vary from trial to trial for different sparsity levels. The yellow region marks experimentally observed levels of sparsity. The inset shows how the mutual information varies for three values of the sparsity *p*, i.e. the fraction of glomeruli that are activated. (C-D) Component separation: ROC curves are shown for three values of *ρ* (color scheme as in panel A) and two relevant values of *p*. Inset: Same curves in semi-log scale.

### Antagonism and olfactory psychophysics

Mixture perception is influenced by interactions throughout the olfactory sensory pathway [6, 51–53]. Evidence for the significant role of receptor-level interactions comes from various direct and indirect measurements [26–28]. Here, we examine the possible relation between the antagonistic effects presented above, and observations for psychophysical experiments on the perception of odor mixtures. The upshot is that the combination of competitive antagonism and masking supports the diverse range of psychophysical effects enumerated hereafter.

Specifically, the list of psychophysical effects relevant here is as follows (see [16,23–25,46]). 1) Inhibition and synergy: The former is the strong reduction of perceived intensities when two odorants are mixed, usually at high concentrations ; synergy is occasionally observed, namely at low concentrations. 2) Masking: When the concentration of a masking agent, such as 2,4,6- trichloroanisole (called cork taint), is increased, the perceived intensity of the odorant decreases. 3) Symmetric and asymmetric suppression: When two odorants of equal perceived intensities are added, they typically suppress each other in a reciprocal fashion so that the perceived intensity of both odorants is still equal but sharply lowered. Asymmetric suppression (sometimes called counteracting), where the intensity of one of the odorant is lowered more than the other, is observed occasionally. 4) Overshadowing: The loss of perception of a less intense odorant when a more intense odorant is present in a mixture.

Results in Fig. 7 illustrate the prevalence of all the aforementioned psychophysical effects as we tune the level of antagonism in the encoding process. We estimated the inferred (or perceived) concentration of a component in the mixture as the concentration at which those precise number of glomeruli corresponding to the odorant would have been activated had the odorant been delivered alone. Fig. 7A demonstrates that inhibition and synergy naturally arise from competitive antagonism. Masking is also readily explained by our model, where the inferred concentration (based on the activated glomeruli) is below the actual concentration as the masking agent’s concentration increases (Fig. 7B). Fig. 7C shows that reciprocal suppression of intense binary mixtures is conspicuously absent for a non-antagonistic model. Suppression is quantified as the fraction of suppressed glomeruli, defined as the fraction of glomeruli which are inactive in the mixture of *A* and *B*, yet they are activated in isolation by odorant *A*(*B*) and not activated in isolation by odorant *B*(*A*). A strong asymmetric suppressive effect is observed when one of the odorants has a capacity for masking.

**Figure 7:**
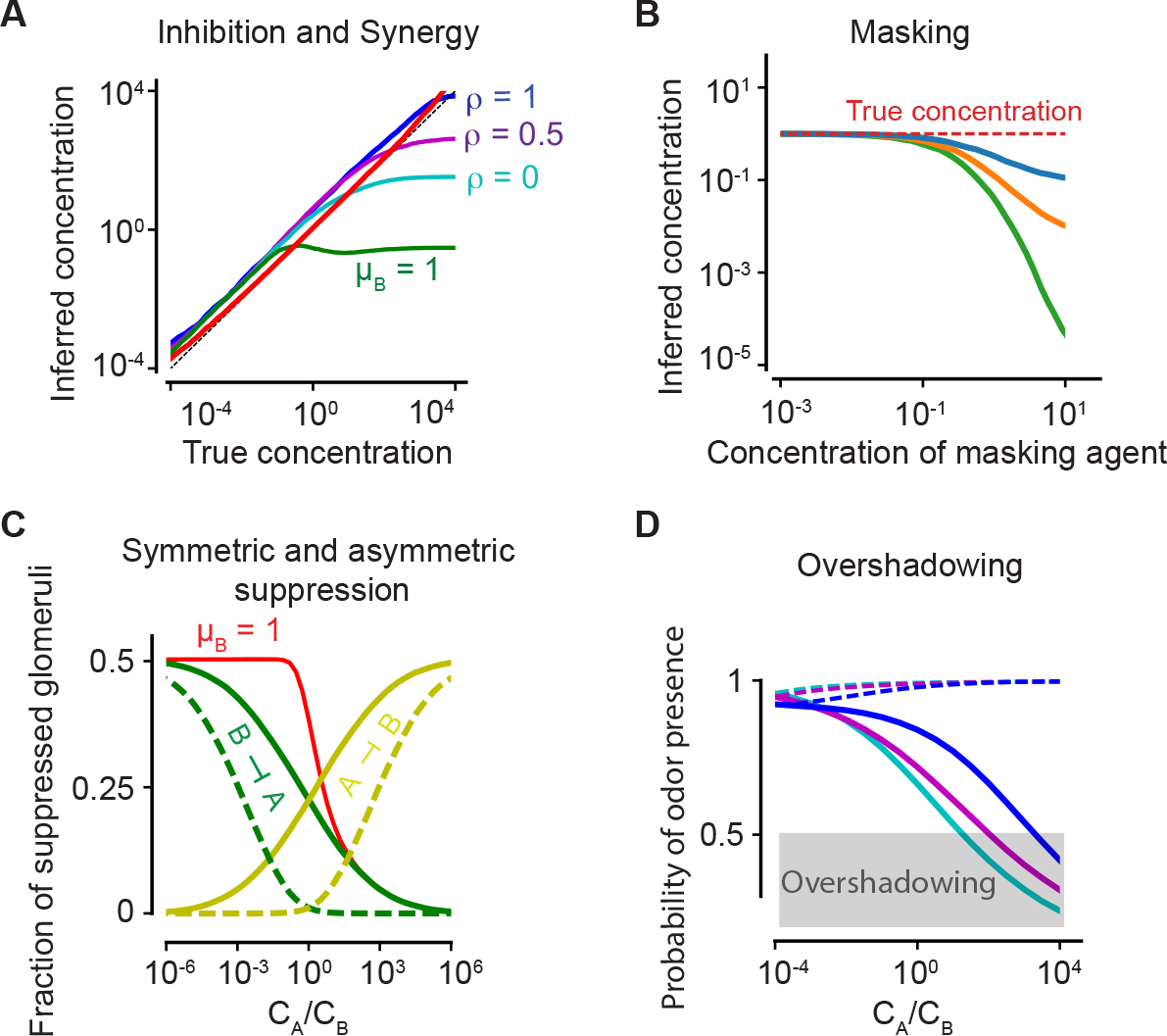
Antagonism and psychophysics. (A) Inhibition and synergy: The inferred *vs* the true concentration for a single odorant *A* (solid, red line) or with an additional odorant *B*. Blue, magenta, cyan: *ρ* = 1,0.5, 0, as defined in (4). Green line: the inferred concentration when *B* has a high masking coefficient *μ*_*B*_, as defined in (19) and (21). In all panels, the sparsity of glomerular activation is *p* = 0.5. (B) Masking: The inferred concentration of A for increasing concentrations of a masking agent B (blue, orange, green: *μ*_*B*_ = 0.4,0.7,1). (C) Symmetric and asymmetric suppression: The fraction of suppressed glomeruli of *A*(*B*) is plotted in green (yellow) against the ratio of concentrations of *A* and *B*. Solid/dashed lines: *ρ* = 0,1. The red line shows the fraction of suppressed glomeruli when *B* also has a propensity for masking. (D) Overshadowing: The probability of presence of *B* as computed by the logistic regressor against the ratio of *A* and *B* concentrations. The dashed lines show the probability of presence of *A*. Color code is as in panel (A).

Finally, to quantify overshadowing, we train a logistic regressor to identify a set of known odorants as in Fig. 4C-D. A weak odorant B is delivered along with a stronger odorant *A* at varying concentration ratios. The probability of presence of *B* as computed by the regressor is compared against the the ratio of concentrations of *A* and *B*. When the probability of presence goes below the detection threshold (set at 0.5), *B* is no longer detected and is “overshadowed”. Fig. 7D demonstrates that overshadowing is intensified by antagonism, in spite of its superior discriminatory performance.

## Discussion

Natural smells are due to mixtures of many chemicals, yet the need for tight stimulus control in experiments often leads to a focus on individual molecular entities. In this paper, we have characterized mixture interactions with a realistic biophysical model and experimentally demonstrated the prevalence of nonlinear mixtures interactions in ORNs of mice. Importantly, we explored how these interactions can naturally lead to “normalization” of the glomerular responses, improve the coding capacity of the olfactory system, and account for many observed perceptual phenomena.

The odorant receptor dynamics in our model is based on a two-step activation process analogous to previous works in vertebrates [36,37] and the fly [54]. In contrast to earlier work, we explicitly model the pathway downstream of receptor activation [34,35], which features the successive steps of cAMP production, allosteric opening of CNG channels, and ultimately current fluxes. This provides a biophysical basis to the cooperativity effects that were previously introduced *ad hoc.* Moreover, this explicit formulation allows us to go beyond pure competitive antagonism, which was reported to explain about half the cases and thus requires generalizations [36]. In particular, non-competitive antagonistic effects, such as our masking model for the non-specific suppression of the cyclic nucleotide-gated channels permit us to account for synergy and inhibition effects that are impossible for competitive antagonism.

A key aspect of two-step activation is that it separates sensitivity of ligand binding from activation efficacy [55]. At the structural level, this distinction is consistent with observations on the binding and activation of GPCRs [56]. The common and parsimonious approximation of a single step (parametrized by a *K_d_* for affinity) in binding models, appears too drastic. Indeed, a signature of multi-step models is different maximum responses to different ligands, in addition to the commonly presented differences in the concentration of half-maximal activation. We found evidence in (ours and previous) experimental datasets for such differences in maximal activation levels.

To assess the nature of mixture interactions in the mouse olfactory system, we performed experiments *in vivo* to mimic natural conditions of the olfactory epithelium and validate our theoretical model. Our motivation for new experiments stemmed from the conflicting evidence for nonlinear interactions in previous datasets ( [36,48,52,57], and the importance of obtaining data from a large number of receptors and over a wide range of concentrations. Indeed, the concentration of odorants was not varied systematically in studies that provided evidence for relatively linear summation of odorant responses to mixtures ( [32,48,57]). The importance of nonlinear interactions visible in our dataset is also shared by insect olfactory receptors ( [51,59]).

Since we imaged responses in populations of ORN axons and terminals, rather than their somata, some caveats need to be pointed out. First, all responses are averages of many ORNs, which could have heterogeneous relation to odorants both due to intrinsic differences [62,63], as well as the access of odorants to the different ORNs [64]. Any heterogeneity is likely to be similar for different odorants, and therefore unlikely to affect the conclusions of our study. Second, measured responses may be influenced by lateral interactions within the olfactory bulb networks, particularly due to GABAb-mediated presynaptic inhibition that can alter ORN calcium signals [60]. However, there is strong evidence that GABAb-mediated reduction of ORN responses are largely intraglomerular [61], which will simply serve as an automatic gain control, hence insensitive to the identity of the odorant. As an additional check, we chose two distinct odorant mixtures with differing extents of overlap in glomerular activation. We found similar mixture effects for both pairs of odorants, which indicates that lateral interactions among glomeruli do not affect our conclusions. A third caveat is that calcium imaging offers only a time-averaged index of ORN activity, and any additional information in timing of spikes may be lost. However, except at the highest concentrations of odorants, several parameters such as latency, number of spikes and average spiking rate appear to be strongly correlated with each other.

A major focus of our work is the functional role of antagonistic interactions. Antagonistic reduction of glomerular activation can be seen as a form of “normalization of activity”. Normalization with increasing stimulus intensity or complexity is common in neural systems [65], and has been thought of as a circuit property that involves inhibitory synaptic interactions [66,67]. In the olfactory system, this was elegantly demonstrated in the *Drosophila* antennal lobe, where activation of increasing number of receptors (or glomeruli) proportionally increases inhibition provided to any one glomerular channel [68]. Similarly, in fish and mouse olfactory bulb, increasing stimulus intensity is thought to recruit populations of interneurons (namely, short axon cells) that inhibit principal cells, leading to blunted activity for higher stimulus intensities. In an extreme example of this, a mouse with a particular receptor forcibly expressed widely has remarkably similar activation of mitral cell population despite the massive increase in the input when the cognate odorant is presented [69].

The key insight from our model is that normalization is granted at the level of receptors by purely statistical reasons, without any additional circuit burden. In the limit of full statistical decorrelation between ligands’ binding affinity and activation efficacy, the distribution of activations across the ORN ensemble for a mixture coincides with that of a single monomolecular odorant, a property which has been confirmed in the fly [70]. Why would we need normalization if the optimal way to preserve information is to simply copy the input signal, i.e. have ORNs functioning as pure relays? Copying, however, requires an unrealistically broad dynamic range, especially for the processing of natural mixtures, where the concentration and the number of components can fluctuate wildly. Normalization at the first layer in the sensory pathway helps avoid early saturation effects that would confound the entire processing pathway. However, nonlinear distortions of the signal do lead to loss of information, and the balance between the two effects calls for their quantification. Our information theoretic calculations demonstrate that detection of a target odorant within a complex mixture is enhanced by antagonistic interactions, and that holds for a wide range of receptor tuning widths, i.e., the average number of activated glomeruli per odorant.

In addition to the above functional advantages, we showed that antagonistic effects are consistent with psychophysical effects observed in mixture perception. Experiments show that the perceived intensity level of an odorant is empirically related to the true concentration of the odorant as a power function, reflecting their proportional relationship on a single logarithmic scale [23,71]. The intensity level for binary mixtures perception is commonly described via a vector sum of the intensities of each component [72-74]. The vector model captures the level independent, symmetric nature of mixture suppression. The biophysical model presented here is consistent with an even broader range of perceptual phenomena, including level independency, synergy, symmetric and asymmetric suppression, masking and overshadowing. The bottomline is that global antagonistic interactions at the ORN level may play a major role in non-trivial perceptual phenomena.

In conclusion, ORNs are far from simple relays, and their strong nonlinear interactions crucially affect olfactory processing. Non-competitive antagonistic mechanisms, such as masking effects discussed here, have not been widely studied in mice and they may only occur for selected odorants. While this experimentally necessitates an extensive experimental dataset, the non-competitive effects presented here make their future investigation particularly relevant. Further, the sensitivity and activation efficacy were inferred indirectly from the mixture response curves of glomeruli, leading to noisy estimates of these values, and motivates a more direct measurement in single ORNs. Finally, the generality of the potential relations highlighted here between ORN antagonism and psychophysical phenomena motivates their exploration in mice, where a broader arsenal of experimental techniques and manipulations can be leveraged.

## Methods

### Experiments

#### Animal Care, General Statements

Male and female olfactory marker protein (OMP)-GCaMP3 mice [43] were used in this study of which hemizygous breeding pairs were maintained in our colony. The age of the animals at the time of the experiments was two to four months. A total of eight animals were used for this study. All the experiments were performed in accordance with the guidelines set by the National Institutes of Health and approved by the Institutional Animal Care and Use Committee at Harvard University.

#### Surgery

Adult mice were anesthetized with ketamine and xylazine (100 and 10 mg/kg, respectively) and the cranial bones over the olfactory bulbs were removed using a 3 mm diameter biopsy punch. The surface of the brain was cleared of debris and a glass coverslip was glued into the vacated cavity in the skull. A custom metal head plate was then affixed to the rear of the skull with dental cement. A shallow well made of dental cement was formed around the coverslip to hold fluid for our water immersion objective.

#### In vivo imaging

Animals were allowed to recover from surgery for at least three days prior to imaging. Prior to each imaging session, animals were anesthetized with an intraperitoneal injection of ketamine and xylazine (90% of dose used for surgery) and their body temperature was maintained at 37°C by a heating pad. A custom-built two-photon microscope was used for in vivo imaging. Fluorophores were excited and imaged with a water immersion objective (20×, 0.95 NA, Olympus) at 920 nm using a Ti:Sapphire laser (Mai Tai HP, Spectra-Physics). Images were acquired at 16-bit resolution and 4 frames/s. The pixel size was 1.218 *μ*m, and fields of view were 365 × 365 *μ*m. The point spread function of the microscope was measured tobe0.51 × 0.48 × 2.12 *μ*m. Image acquisition and scanning were controlled by custom-written software in Labview [75]. Emitted light was routed through two dichroic mirrors (680dcxr, Chroma and FF555-Di02, Semrock) and collected by a photomultiplier tube (R3896, Hamamatsu) using filters in the 500-550 nm range (FF01-525/50, Semrock). Signals were digitally filtered prior to acquisition (Stanford Research Systems).

#### Odorant stimulation

Monomolecular odorants (Penta Manufacturing) were used as stimuli and delivered by a custom-built 16-channel olfactometer controlled by custom-written software in Labview (National Instruments). The olfactometer was outfitted with two separate odorant pairs, ethyl valerate/allyl butyrate and methyl tiglate/isobutyl propionate. The initial concentration series for each odorant was 80%, 16%, 8%, 1.6%, 0.8%, 0.16%, 0.08% (v/v) in mineral oil and further diluted 16 times with air. To generate the highest final odorant concentrations, two valves containing 80% odorant were opened concurrently to reach a total final concentration of 10%. odorants were presented for 2s to prevent adaptation at the strongest concentrations. Clean air was delivered through the inlet at a constant flow both preceding and following odorant delivery and all mixtures were generated by air-phase dilutions just prior to reaching the animals' nose to prevent the formation of secondary or tertiary odorant compounds. For all experiments, odorants were delivered 3-5 times each with an inter-stimulus interval of at least 40s.

Final odorant concentrations for each component and mixture were verified using a photoionization detector (PID; Aurora Scientific). The PID inlet was placed in the same spatial location as the animals nose to assay the odorant concentration experienced by the animal. Each odorant dilution series and mixture was normalized to the largest PID measurement.

#### Data extraction

The response of individual glomeruli was measured by manually drawing ROI boundaries on maximum intensity projection images. To temporally measure fluorescence intensity, the mean intensity of all pixels within each ROI boundary was calculated for each frame. Data were corrected for photobleaching by subtracting blank trials. The baseline fluorescence was established by calculating the mean intensity in the 20 frames preceding odorant stimulation.

#### Data analysis

The time series of the response of each glomerulus for each trial, measured as the relative change in fluorescence levels, ΔF/F, was first smoothed using an exponential smoothening filter with a smoothening parameter of 5 frames (1.25 s). The peak of the smoothened response curve was averaged over at least three trials of the same odorant stimulus to obtain the mean peak response for different concentrations of the odorants and their mixtures. A glomerulus was removed if it did not respond to any of the odorants, where lack of response was declared if the peak response was below a threshold set at three times the standard deviation of the fluorescence level 20s before odorant stimulation. The parameters *κ* and *η* for the odorants in the pair were determined by simultaneously fitting the mean peak response from the individual odorants and the mixture using equations (2) and (3). The solid red, blue and magenta lines in Fig. 3 were obtained by using the best fit parameters in equation (2) for each odorant and in equation (3) for the mixture. The dashed magenta line in Fig. 3 was obtained from a sigmoidal fit to only the mixture data points.

### Modeling

#### Competitive binding

When a mixture of *K* monomolecular odorants *X*_1_, *X*_2_,…, *X_k_* at concentrations *C*_1_, *C*_2_,…,*C*_*K*_ is presented to an ORN, the odorants compete for the finite number of receptors available on the olfactory cilia. In this Section, we derive the response of the ORN in such a case of competitive binding under the assumptions stated below.

The binding of an odorant to a receptor induces the activation of the odor-receptor complex, via a two-step GTP-mediated phosphorylation process [34]. For a mixture of odorants, the binding dynamics reads:

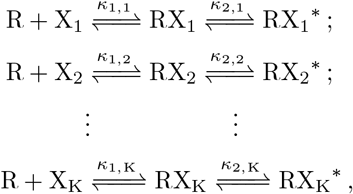

where *RX*_*i*_ and 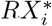 symbolize the bound and activated complexes (i = 1, 2,…, *K*), while *R*, *B*_*i*_ and *A*_*i*_ denote the number of unbound receptors, receptors bound by odorant *i* but inactive, and receptors activated by odorant *i*, respectively. The concentration *C*_*i*_ of the various odorants is supposed to be in excess for the total number of receptors 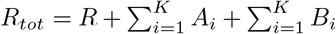. The parameters *κ*_1,*i*_ and *κ*_2,*i*_ are the ratio of the backward rates to the forward rates for the two reaction steps involving odorant *i*. We introduce only the ratio of the rates because the time scale of the slowest step, the activation of the odor-receptor complex, is estimated to be a few hundred milliseconds [40]. For delivery times of a few seconds or longer, we can then assume equilibrium. By using the steady-state relations *B*_*i*_ = *C*_*i*_*R*/*κ*_1,*i*_, *A*_*i*_ = *B*_*i*_/*κ*_2,*i*_, and the above equation for *R*_*tot*_, we obtain that the number of activated receptors bound to odorant *i* at equilibrium is

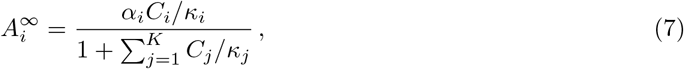

with 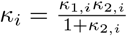 and 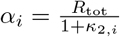 .

A chain of steps follows receptors’ activation in the transduction pathway. First, activated receptors convert ATP into cAMP molecules via adenylyl cyclase III. Then, the cAMP molecules diffuse locally and open nearby cyclic-nucleotide-gated (CNG) ion channels. The open CNG channels are permeable to Ca^2+^ (and Na^+^) ions, which are crucial in regulating further downstream processes and for adaptation [42]. Finally, Ca^2+^ ions bind to calmodulin (CaM) and the formed complex (Ca-CaM) inhibits the CNG channels, leading to adaptation and, possibly, the termination of the response. The primary depolarizing current is carried by an Cl^−^ efflux out of the cell through Ca^2+^ regulated Cl^−^ channels. We now proceed to model these various steps.

First, since CNG channels are spread out along the cilia membrane and cAMP diffusion is restricted to the site of its production [77,78], successful receptor activation events are largely independent. Indeed, the electrical response is consistent with Poisson statistics, and the voltage-clamped current response close to the threshold is linear [76]. At concentrations much larger than the threshold, the production of cAMP is linearly proportional to the number of activated receptors, as evidenced by the linear increase of the rate of production of cAMP with time. The degradation of cAMP occurs on a single time scale of a few hundred milliseconds [78]. The effective cAMP dynamics is then succinctly written as:

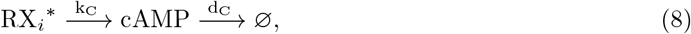

where *κ*_*C*_ is the rate of production of cAMP (and implicitly includes the concentration of the converted ATP, which is supposed to be in excess and thereby treated as fixed), *d*_*C*_ is the rate of degradation, and the index *i* runs over the set of *K* rate equations. Since the production of cAMP occurs immediately downstream of activation and is independent of the activating odorant, the rate of production is simply proportional to the total number of activated receptors. If *C* is the intracellular cAMP concentration, we conclude that the steady state cAMP concentration *C*^∞^ is

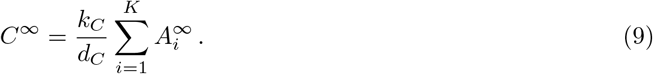

Second, CNG channels have four binding sites for cAMP and exhibit allosteric cooperativity [79], which is generally represented as [80] :

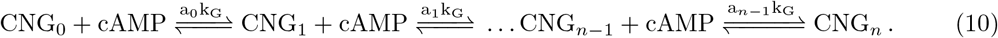

Here, *k*_*G*_ is an overall rate, *n* is the number of allosteric binding sites for cAMP, and *a*_0_, *a*_1_,…, a_*n*−1_ modulate the various steps of the reactions. For allosteric cooperativity, *a*_0_ < *a*_1_, *a*_2_,…, which reflects the fact that the binding of one cAMP molecule promotes the binding of further cAMP molecules. Here, we have ignored the inhibitory effect of Ca-CaM, which will be introduced below. Allosteric cooperativity leads to response functions of the Hill form [80]. In the limit of strong cooperativity, most of the CNG channels are expected to be either unbound or fully bound, and the steady state number of fully bound CNG channels reduces then to

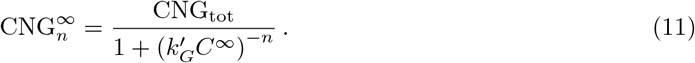

Here, *C*^∞^ is the expression (9), 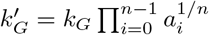 and CNG_to_t is the total number of CNG channels. The Ca^2+^ current is directly proportional to the number of fully bound CNG channels and decreases at a constant rate [39] :

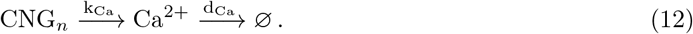

Third, the production of the CaCaM complex by Ca^2+^ and CaM is described as:

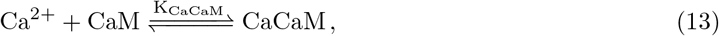

where *K*_*CaCaM*_ is the ratio of the forward and backward rates. The effect of calmodulin-mediated feedback inhibition is accounted by assuming CaCaM modulates the CNG opening rate as 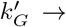 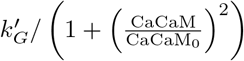. The previous form is empirical, yet we verified that its precise shape and the alue of the Hill coefficient do not modify numerical results below as long as a steep sigmoidal shape lolds. With the values of the parameters used in our model (Table S1), CaCaM acts on the CNG channel before the cAMP and the activated receptors reach steady state and terminates the response. The resulting set of differential equations does not lend to analytical treatment but can be numerically integrated to give a time series of the ORN response, as shown in Fig. 1B. Numerical curves indicate data not shown) that the Ca^2+^ peak response terminated by CaM is roughly proportional to the steady state Ca^2+^ response without CaM, i.e. 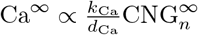, which is the approximation that we shall use hereafter.

Finally, the Cl^−^ current is the predominant component of the currents that depolarize ORNs [39,58] rnd is mediated by Ca^2+^-gated Cl^−^ ion channels. This current response induced due to Ca^2+^ is again a Hill function of coefficient greater than one, suggesting further cooperativity [81]. We can formally write the steady state Cl^−^ current as 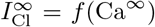, where *f* is some unknown function. The firing rate response *F* of the ORN is assumed to be proportional to the current, so that 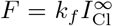, where f is a constant.

By combining all the equations above, we can write the firing rate as a function of the odorant concentrations *C*_*i*_ :

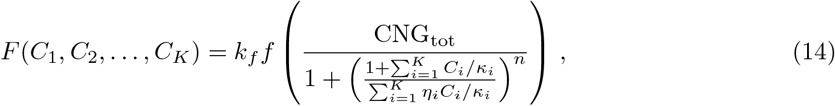

where the odorant-receptor dependent parameters for the *i*th odorant are written explicitly:

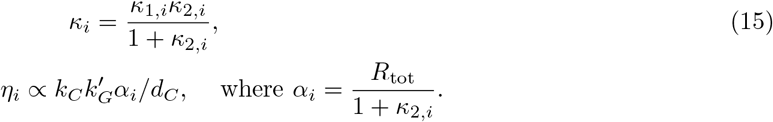

As mentioned in the main text, for each odorant *i*, *κ*_*i*_ and *η*_*i*_ carry the dependence on the odorants and control their interactions within mixtures. Conversely, the unknown function *f* is related to downstream processes and therefore expected to not depend on the odorant and receptor type. This point, together with the reduction in the number of free parameters, was our rationale for using the approximation of a linear function *f*. Then, the proportionality constant in *f* is lumped together with *k*_*f*_ and *CNG*_*tot*_ in (14) into a single parameter *F*_max_, which defines the maximum physiologically possible firing rate of the neuron. Both *F*_max_ and *n* are constants that do not depend on the odorant or the receptor type.

We can finally write a general expression for the ORN response to a mixture of *K* odorants with concentrations *C*_1_, *C*_2_,…, *C*_K_. Denoting the total concentration by 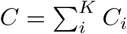 and the contribution of the *i*th component by *β*_*i*_ = *C*_*i*_/*C*, it follows from (14) that the response reads

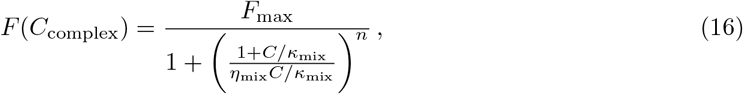

where the “effective” mixture parameters *η*_mix_ and *κ*_mix_ are

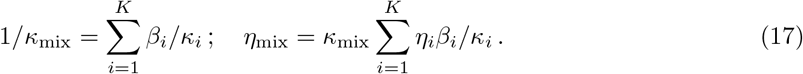

A complex odorant can therefore be treated in a manner similar to monomolecular odorants, namely, by specifying its effective sensitivity and activation efficacy to each receptor type. Note that this holds generally true, irrespective of the linear *f* chosen in (14) to limit the number of free parameters.

The parameters used in Fig. 1 are as follows - we first define *k*_1_, *k*_−1_ as the forward and backward rates for the binding step of (1), and *k*_2_,*k*_−2_ as the forward and backward rates for the activation step. For panels B,C and D, we use *k*_1_ = 100*s*^−1^, *k*_−1_ = 100*s*^−1^, *k*_2_ = 2*s*^−1^, *k*_−2_ = 2*s*^−1^ for odorant A and *k*_1_ = 80*s*^−1^, *k*_−1_ = 100*s*^−1^, *k*_2_ = 0.4*s*^−1^, *k*_−2_ = 2*s*^−1^ for odorant B. The concentration is unity in panels B, F and G. For panels E, F and G, the parameters for odorant A are used. In all panels, we use *k*_*C*_ = 2*s*^−1^, *d*_*C*_ = 1*s*^−1^, 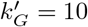, CNG_tot_ = 1, *n* = 4, *k*_*Ca*_ = 20*s*^−1^, *d*_*Ca*_ = 0.5*s*^−1^, *k*_*CaCaM*_ = 1*s*^−1^, CaCaM_0_ = 0.05.

#### Masking

Here, we present a phenomenological description of non-competitive masking processes.

We suppose that masking agents bind sites on the lipid bilayer and compete for their limited number. The suppression timescale and off-timescale are smaller than a few hundred milliseconds [22], justifying the assumption of steady state. In steady state, the occupancy fraction of the ith masking agent with concentration *M*_*i*_ and binding affinity 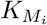 is

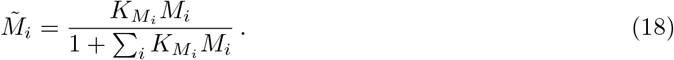

The disruption of a CNG channel conformation due to agent *i* is supposed to alter the affinity of cAMP to one of its binding sites on the channel in the reactions (10). The energy of the cAMP bound state is increased by Δ*ϵ*_*i*_ and its probability is reduced by the corresponding Gibbs factor 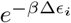, where *β* = 1/*kT* is the inverse temperature. The resulting reduction in the opening of the channels is most conveniently accounted for by a mean-field approach where the channel opening rate 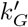 appearing in (11) is modified by the masking agents. In other words, 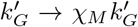 with the suppression factor χ*M* < 1 derived below. It follows from the definition (14) of *η* that a modification of 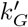 by *χM* carries over to *η* as *η* → *χMη*. Therefore, when saturating concentrations of excitatory odorants are presented together with masking agents that produce a masking coefficient *χM*, the maximal firing rate is reduced as

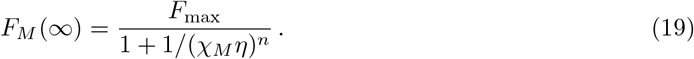

which reflects the masking effect.

The dependence of the suppression factor *χM* on the concentrations *M*_*i*_ of the masking agents is estimated as follows. Let us denote the radius of disruption of the channels by a single masking molecule on the lipid bilayer by *r*, and the surface density of masking binding sites by *σ*. The typical number of masking binding sites surrounding a given CNG channel is then λ = *πσr*^2^. The number *n*_mask_ of masking binding sites within distance *r* of a CNG channel is assumed to be Poisson distributed, i.e. 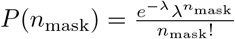. For a given number of sites *n*_mask_, the vector of their occupancy numbers ***I*** = (*i*_l_, *i*_2_, ․.*i*_*K*+1_) is distributed following a multinomial distribution with probabilities given by (18), i.e. 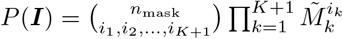. The index *K* + 1 corresponds to unoccupancy, 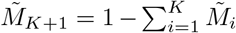 and *i*_*K*+1_ = *n*_mask_ − ∑_*k*_ *i*_*k*_. The probability of each ***I*** is proportional to its Gibbs factor *e*^−*β*Δ*ϵ*(***I***)^, where Δ*ϵ*(***I***) is the energy shift to the binding of masking agents.

We consider the first step in (10); similar arguments hold for successive ones. The unmodified *a*_0_*k*_*G*_ = *e*^−*βϵ*^ is the ratio between the probability for a channel to be cAMP bound or cAMP unbound, and e is their energy difference. In the presence of masking, there are multiple cAMP bound and unbound states, which differ in their occupancy of the masking binding sites. The sum over all those states defines the probabilities *P*_*b*_ and *P*_*u*_ of cAMP bound and unbound, respectively. The suppression factor *χM* that modifies 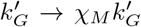 is obtained as the ratio (*e*^*βϵ*^*P*_*B*_/*P*_*u*_)^l/*n*^, where the 1/*n* power stems from the definition of 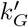 in (11). The sum *P*_*b*_ is obtained by combining all the previous factors:

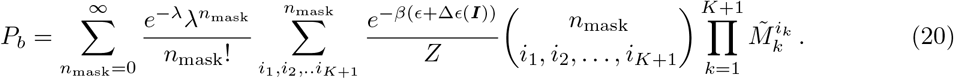

where *Z* is a normalization factor. Assuming the masking sites do not affect the energy of the channels when cAMP is unbound, the sum *P*_*u*_ has a similar expression with *ϵ* + Δ*ϵ* = 0. It is then verified that *P*_*u*_ = 1/*Z*. As for *P*_*b*_, the simplest possible assumptions are that 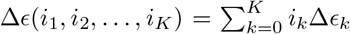 is additive, and the masking binding sites are dilute, i.e. λ is small. Eq. (20) reduces then to

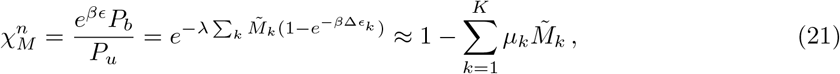

where 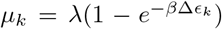 satisfy 0 ≤ *μ*_*k*_ ≤ 1, and the same inequality holds for *χM*. In general, masking agents can affect multiple CNG channel subunits [82]. If a masking agent affects the binding of cAMP to *j* CNG subunits, the suppression effect is *χM* = (1 − ∑_*i*_*μ*_*i*_*M̃*_*i*_)^*m*^ with *m* = *j/n*.

The ratio 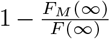 in (19) is plotted in Figure 4B and compared to experimental data. In Figure 4C,D, the parameters for generating the response curves for odorants A and B are *κ*_*A*_ = 1, *κ*_*B*_ = 1, *η*_*A*_ = 1,*η*_*B*_ = 5. For synergy (Fig. 4C), the masking parameters are *K*_*M,A*_ = 10^−5^,*K*_*M,B*_ = 10^−1^,*μ*_*A*_ = 0,*μ*_*B*_ = 0.7, while the corresponding parameters for inhibition (Fig. 4D) are *K*_*M,A*_ = 1,*K*_*M,B*_ = 10 ^−5^, *μ*_*A*_ = 0, *μ*_*B*_ = 0.7. The parameter *m* is chosen to be unity in both cases.

#### Olfactory encoding model

Every odorant is defined by a vector of binding sensitivities ***κ*^−1^** and a vector of activation efficacies *η*, each with dimensionality *N*, where *N* = 250 is the chosen number of receptor types. An odorant's binding sensitivity to a particular receptor type is drawn independently from a log-normal distribution 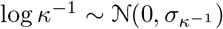, where its standard deviation 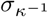 is set to 4 to obtain a six orders of magnitude separation between the most sensitive and least sensitive receptor types [47]. The activation efficacies are similarly drawn independently for each receptor type such that 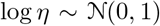. The measure of antagonism, *ρ*, is defined as the Pearson correlation coefficient between log *κ*^−1^ and log *η*:

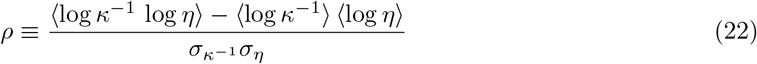

where *σ*^2^ denotes the variance of the random variables and the angular brackets denote expectation values. To generate an odorant-receptor pair, first log *η* is drawn from the standard normal distribution. Then, log *κ*^−1^ is generated with correlation *ρ* as 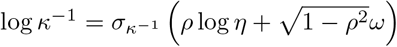, where *ω* is drawn from a standard normal distribution. At saturating concentrations of an odorant, the peak firing rate it elicits for a receptor type with activation efficacy *η* is 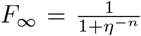 (see (16), where *F*_max_ can be chosen to be unity). The rescaled glomerular activation vector ***y*** (each component is rescaled between 0 and 1) is given by 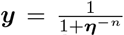, where the transformation is performed on each component of the vector. The probability *p* that each component exceed a threshold τ is given by the probability that a random variable drawn from a standard normal distribution exceed 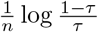. This probability *p* represents the sparsity of the glomerular activations ***z*** at saturating concentrations after thresholding. The sparsity is set by selecting 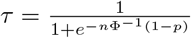, where Φ is the cumulative distribution function for
a standard normal random variable.

### Figure-ground segregation and component separation

To quantify the performance of the encoding model in figure-ground segregation, we compute the mutual information between the absence (*T* = 0) or presence (*T* = 1) of the target odorant *T* and the glomerular activation pattern ***z***. Noise is introduced due to the presence of background odorants of unknown sensitivities and activation efficacies. The mutual information controls the performance of an optimal Bayesian decoder in detecting a target odorant in a background by using the glomerular activation pattern as input. The mutual information is defined as

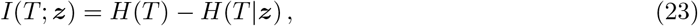

where *H*(*T*) is the entropy of target presence or absence, which equals one bit (since the target is present in half the trials). The second term on the right hand side *H*(*T*|***z***) (in bits) is given by

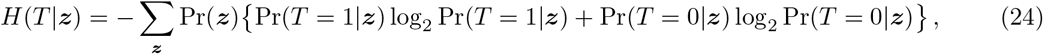

where Bayes’ formula yields 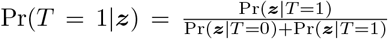, and 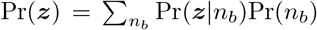, where Pr(*n*_*b*_) is the distribution of the number of background odorants. *H*(*T*|***z***) is estimated numerically by using Monte Carlo sampling. The quantities Pr(***z***|*T* = 1) and Pr(***z***|*T* = 0) are also computed numerically by observing that 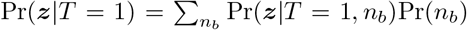. Due to the independence of the receptor types and since the background odorants are independently drawn, the probability Pr(***z***|*T* = 1, *n*_*b*_) factorizes into *N* multiplicative terms, each of which can be pre-computed prior to Monte Carlo sampling. When the number of background odorants fluctuate (as in Fig. 6B), we draw n_*B*_ from a truncated exponential distribution with a mean of 32 and truncated at a maximum of 128 background odorants.

For the component separation task, we use an ensemble of linear classifiers as our decoders. A linear classifier computes the probability of presence of an odorant from the glomerular activation pattern ***z*** as 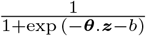, where ***θ*** and *b* are the vector of learned weights and bias, respectively. First, linear classifiers are trained to identify odorants from a fixed set 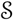 of 500 odorants. During the training phase, each classifier is trained to identify the presence of its target against a background of one to ten odorants also chosen from 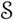. In each trial of the test phase, 1 to 20 odorants are uniformly chosen from 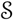 and the component separation performance is measured using the fraction of correct identifications (hit rate) and the number of false positives (FPs). An odorant is declared to be present if the probability of presence exceeds a detection threshold. The hit rate and the number of false positives are modulated by sliding the detection threshold of each linear classifier, yielding the generalized ROC curves in Fig. 6C,D.

### Statistics

Statistical significance was determined using nonparametric Wilcoxon rank-sum test.

### Data availability

All data sets of ORN calcium signals are available upon request.

### Code availability

Code for the modeling can be accessed at: https://github.com/greddy992/Odor-mixtures

## Acknowledgments

We are grateful to JP Rospars for sharing the experimental dataset from Ref. [36]. We also thank Vikrant Kapoor for technical assistance and members of the Murthy Lab for helpful discussions. GR and MV were partially supported by the Simons Foundation Grant 340106. The experimental work was partly supported by a grant from the NIH (R01 DC014453) and JZ was supported by NIH Fellowship F32 DC015938.

---

**Figure Supplement 1:**
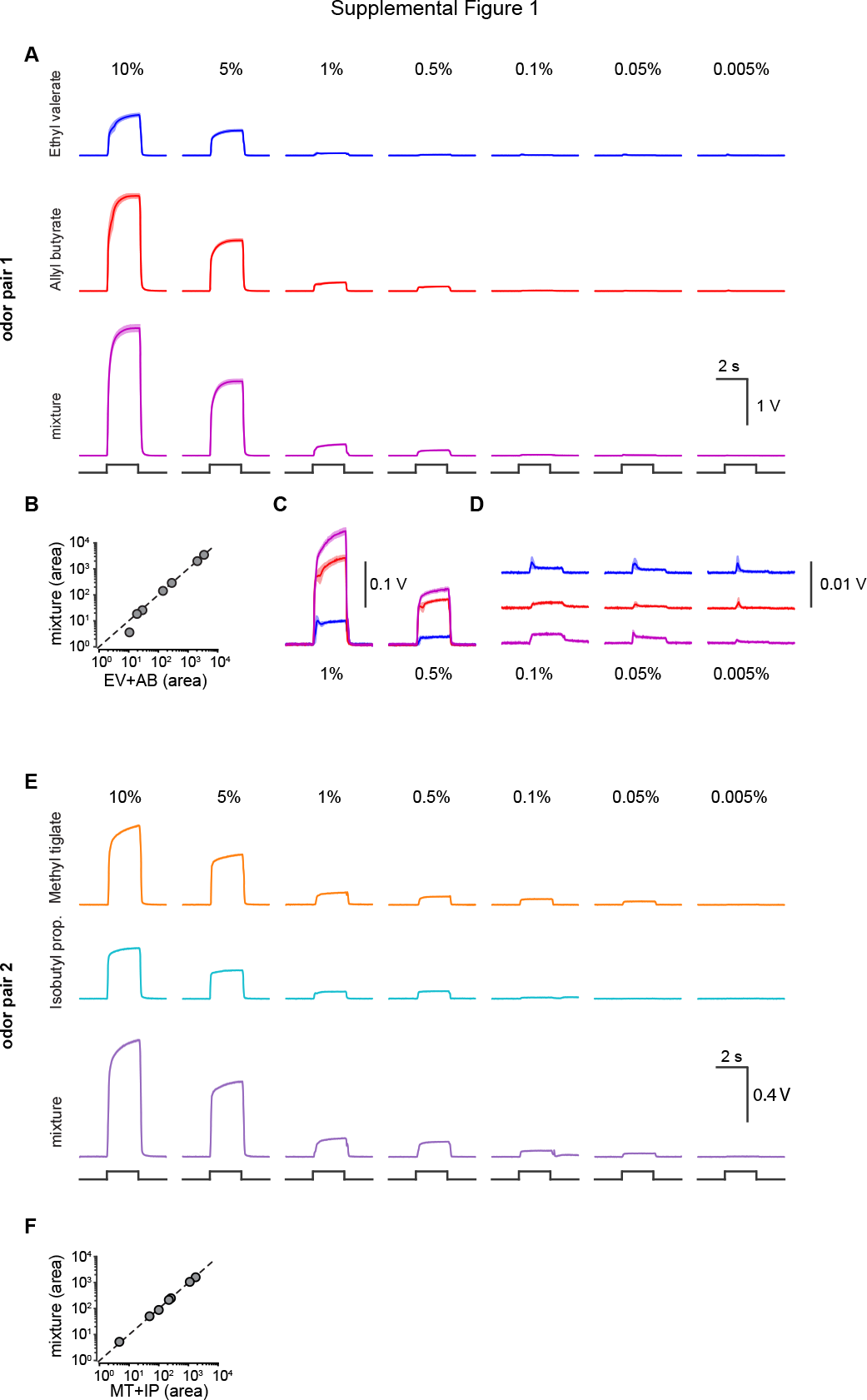
Photoionization detector measurements of odor concentrations. A. Mean photoionization detector (PID) signals from five repeats for both odors in pair 1 and an equiproportionate mixture at each concentration. Shaded area is the s.e.m. Black line below indicates the digital signal used to open each solenoid. B. Scatter plot of the area under the PID signal measured when the solenoid was open for each odor concentration. X-axis is the sum of the area under both odors and y-axis is the area under the mixture response. C. Expanded and overlaid PID signals from odors measured at 1% and 0.5%. D. Expanded PID signals from odors at 0.1%, 0.05%, and 0.005%. Signals are expanded 100 times from part A. E. Same as in part A for odor pair 2. F. Same as in B for odor pair 2.

**Figure Supplement 2:**
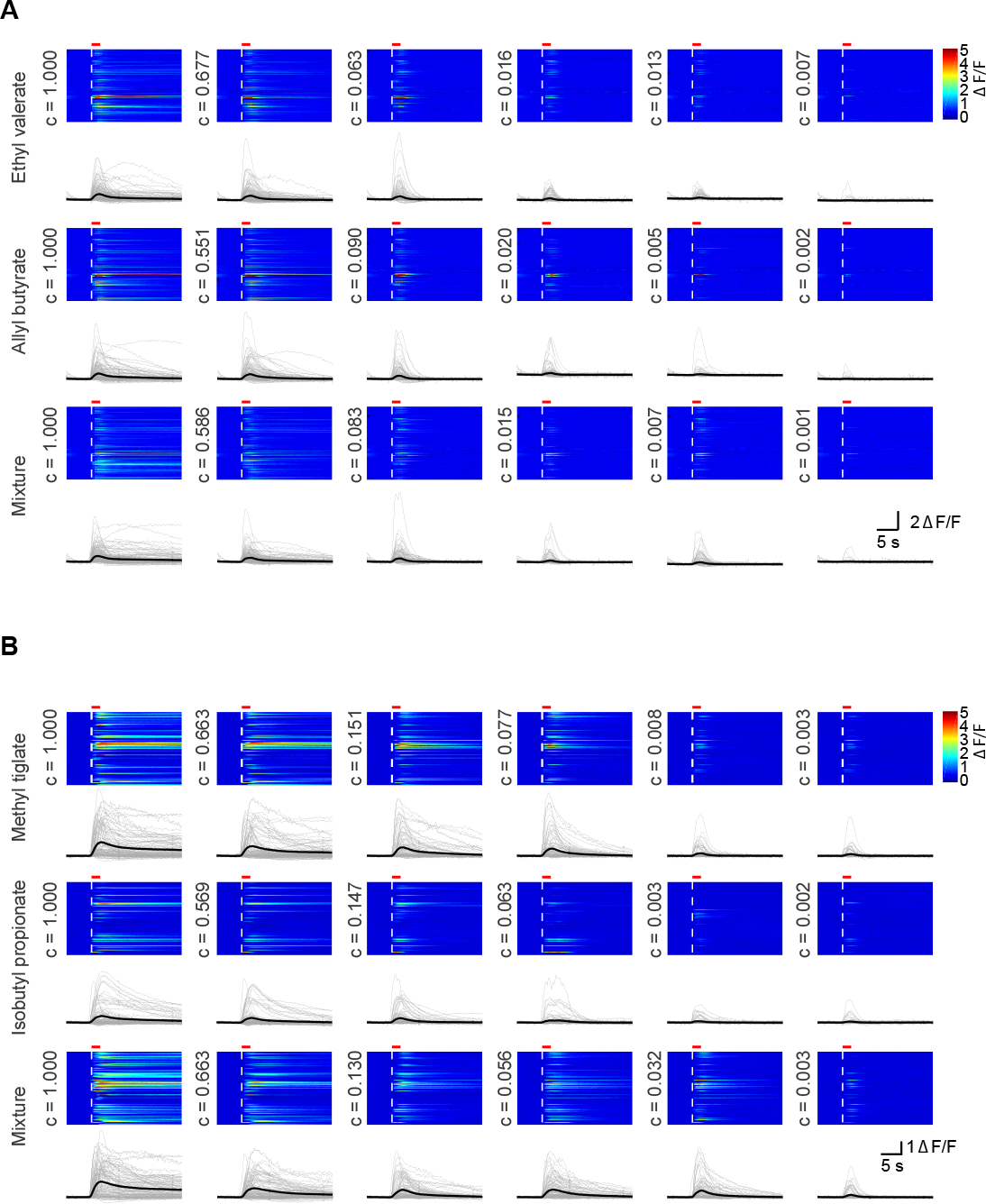
ORN calcium signals from all glomeruli. A. Calcium signals from glomeruli used for odor pair 1 at each odor concentration. Each row in the heat map is one glomerulus and its index is maintained for all odors and concentrations. Dashed white line in heat maps indicates the odor onset and the solid red bar indicates the odor duration. Below each heat map individual glomerular traces are plotted in grey and the mean across all glomeruli is the thick black line. Normalized odor concentration is indicated to the left of each plot. B. Same as in part A for odor pair 2.

## References

[1] B. W. Ache, A.M. Hein, Y.V. Bobkov, J.C. Principe, Smelling Time: A Neural Basis for Olfactory Scene Analysis, Trends in Neurosciences, 39(10):649–655, 2016.

[2] R. T. Cardé and M. A. Willis, Navigational Strategies Used by Insects to Find Distant, Wind-Borne Sources of Odor, Journal of Chemical Ecology, 34(7):854–66, 2008.

[3] J. A. Gottfried, Central Mechanisms of Odour Object Perception, Nature Reviews Neuroscience, 11(9):628–41, 2010.

[4] J. J. Hopfield, Odor space and olfactory processing: Collective algorithms and neural implementation, Proc. Natl. Acad. Sci, 96(22):12506–12511, 1999.

[5] J. D. Howard, and J. A. Gottfried, Configural and Elemental Coding of Natural Odor Mixture Components in the Human Brain, Neuron, 84(4):857–69, 2014.

[6] A. Jinks, D. G. Laing, A limit in the processing of components in odour mixtures, Perception, 28(3):395–404, 1999.

[7] J. T. Knudsen, L. Tollsten, and L. G. Bergstrom, Floral Scents-a Checklist of Volatile Compounds Isolated by Head-Space Techniques, Phytochemistry, 33(2):253–280, 1993.

[8] M. Pentzek, B. Grass-Kapanke and R. Ihl, Odor Identification in Alzheimer's Disease and Depression, Aging Clin Exp Res, 19:255–58, 2007.

[9] R. A. Raguso, Wake Up and Smell the Roses: The Ecology and Evolution of Floral Scent, Annual Review of Ecology, Evolution, and Systematics, 39(1):549–69, 2008.

[10] J. A. Riffell, Olfactory Ecology and the Processing of Complex Mixtures, Current Opinion in Neurobiology, 22(2):236–42, 2012.

[11] J. A. Riffell, L. Abrell and J. G. Hildebrand, Physical Processes and Real-Time Chemical Measurement of the Insect Olfactory Environment, Journal of Chemical Ecology, 34(7):837–53, 2008.

[12] J. A. Riffell, E. Shlizerman, E. Sanders, L. Abrell, B. Medina, J. A. Hinterwirth, and J. N. Kutz, Flower Discrimination by Pollinators in a Dynamic Chemical Environment, Science, 344(6191):1515–18, 2014.

[13] D. Rokni, V. Hemmelder, V. Kapoor and V. Murthy, An olfactory cocktail party: figure-ground segregation of odorants in rodents, Nature Neuroscience, 17:1225–1232, 2014.

[14] R. J. Stevenson and D. A. Wilson, Odour Perception: An Object-Recognition Approach, Perception, 36(12):1821–33, 2007.

[15] P. Szyszka and J. S. Stierle, Mixture Processing and Odor-Object Segregation in Insects, Progress in Brain Research, 208:63–85, 2014.

[16] T. Thomas-Danguin, et al, The perception of odor objects in everyday life: a review on the processing of odor mixtures, Frontiers in Psychology, Vol. 5, No. 504, 2014.

[17] AK Abbas, AH Lichtman, S Pillai, Cellular and Molecular Immunology, Saunders Elsevier Philadelphia, 2010.

[18] P. Francois et al, Phenotypic model for early T-cell activation displaying sensitivity, specificity, and antagonism, Proc. Natl. Acad. Sci., 110(10), E888–E897, 2013.

[19] J.B. Lalanne and P. Francois, Chemodetection in fluctuating environments: receptor coupling, buffering, and antagonism, Proc. Natl. Acad. Sci., 112(6):1898–903, 2015.

[20] Y. Oka, M. Omura, H. Kataoka and K. Touhara, Olfactory receptor antagonism between odorants, The EMBO Journal, 23(1):120–126, 2004.

[21] H. Takeuchi, H. Ishida, S. Hikichi and T. Kurahashi, Mechanism of olfactory masking in the sensory cilia, J. Gen Physiology, 133(6):583–601, 2009.

[22] T. Kurahashi, G. Lowe and A. Menini, Suppression of Odorant Responses by Odorants in Olfactory Receptor Cells, Science, Vol. 265, pp 118–120, 1994.

[23] H. T. Lawless, Olfactory Psychophysics, In Tasting and Smelling, 1st ed., edited by G. K. Beauchamp and L. Bartoshuk, Academic Press, 1997.

[24] A. Keller and L. Vosshall, Human Olfactory Psychophysics, Current Biology, Vol. 14, No. 20, 2004.

[25] Doty, R.L. and Laing, D.G, Psychophysical Measurement of Human Olfactory Function, Including Odorant Mixture Assessment. In Handbook of Olfaction and Gustation 2nd Edition, edited by Doty, R.L., Marcel Dekker, New York, 2003.

[26] G. A. Bell, D. G. Laing and H. Panhuber, Odour mixture suppression: evidence for a peripheral mechanism in human and rat, Brain Research, 426:8–18, 1987.

[27] D. G. Laing and M. E. Willcox, An investigation of the mechanisms of odor suppression using physical and dichorhinic mixtures, Behavioural Brain Research, 26:79–87, 1987.

[28] M. A. Chaput et al, Interactions of odorants with olfactory receptors and receptor neurons match the perceptual dynamics observed for woody and fruity odorant mixtures, European Journal of Neuroscience, 35:584–597, 2012.

[29] A. Koulakov, A. Gelperin and D. Rinberg, Olfactory Coding With All-or-Nothing Glomeruli, J Neurophysiol, 98:3134–3142, 2007.

[30] Y. Zhang and T. Sharpee, A Robust Feedforward Model of the Olfactory System, PLoS Comput Biol 12(4): e1004850, 2016.

[31] D. Zwicker, A. Murugan and M. Brenner, Receptor arrays optimized for natural odor statistics, Proc. Natl. Acad. Sci, 113(20):5570–5575, 2016.

[32] Mathis et al., Reading Out Olfactory Receptors: Feedforward Circuits Detect Odors in Mixtures without Demixing, Neuron, 91, 1110–1123, 2016.

[33] A. Grabska-Barwinska et al, A probabilistic approach to demixing odors, Nature Neuroscience, 20, 98–106, 2017.

[34] S. Pifferi, A. Menini and T. Kurahashi, Signal Transduction in Vertebrate Olfactory Cilia, In The Neurobiology of Olfaction, edited by A. Menini, CRC Press/Taylor&Francis, 2010.

[35] S. J. Kleene, The Electrochemical Basis of Odor Transduction in Vertebrate Olfactory Cilia, Chem. Senses, 33:839–859, 2008.

[36] J.-P. Rospars et al, Competitive and Noncompetitive Odorant Interactions in the Early Neural Coding of Odorant Mixtures, Jour. of Neuroscience, 28(10):2659–2666, 2008.

[37] G. Cruz and G. Lowe, Neural coding of binary mixtures in a structurally related odorant pair, Scientific Reports, 3: 1220, 2013.

[38] A. Mamman, J.P. Simpson, A. Nighorn, Y. Imanishi, K. Palezewski, G.V. Ronnett, C. Moon. Hippocalcin in the olfactory epithelium: a mediator of second messenger signaling, Biochemical and Biophysical Research Communications, 322(14):1131–1139, 2004.

[39] A. Boccaccio, A. Menini, Temporal development of cyclic nucleotide-gated and Ca^2+^-activated Cl^−^ currents in isolated mouse olfactory sensory neurons, J Neurophysiol., 98:153–60, 2007.

[40] J. P. Rospars, P. Lansky, A. Duchamp, P Duchamp-Viret, Relation between stimulus and response in frog olfactory receptor neurons in vivo, European Journal of Science, 18(5):1135–54, 2003.

[41] H. Takeuchi and T. Kurahashi, Olfactory Transduction Channels and Their Modulation by Varieties of Volatile Substances, In Chapter 10 of Taste and Smell, edited by D. Krautwurst, Vol. 23, pp 115–149, 2016.

[42] T. Kurahashi and A. Menini, Mechanism of odorant adaptation in the olfactory receptor cell, Nature, 385:725–29, 1997.

[43] Isogai, Y., Si, S., Pont-Lezica, L., Tan, T., Kapoor, V., Murthy, V. N., and Dulac, C. (2011). Molecular organization of vomeronasal chemoreception. Nature, 478(7368), 241–245, 2011.

[44] A. Marasco, A. D. Paris and M. Migliore, Predicting the response of olfactory sensory neurons to odor mixtures from single odorant response, Scientific Reports, 6, No. 24091, 2016.

[45] Ukhanov et al, Inhibitory Odorant Signaling in Mammalian Olfactory Receptor Neurons, J Neurophysiol, 103:1114–1122, 2010.

[46] H. Takeuchi, H. Kato, T. Kurahashi, 2,4,6-trichloroanisole is a potent suppressor of olfactory signal transduction, Proc. Natl. Acad. Sci., 110(40):16235–16240, 2013.

[47] H. Saito et al, Odor Coding by a Mammalian Receptor Repertoire, Sci. Signaling, 2(60):ra9, 2009.

[48] D. Y. Lin, S. D. Shea and L. Katz, Representation of Natural Stimuli in the Rodent Main Olfactory Bulb, Neuron, 50, 937–949, 2006.

[49] E.R. Soucy et al, Precision and diversity in an odor map on the olfactory bulb, Nature Neuroscience 12:210–220 (2009).

[50] R. Vincis et al, Dense representation of natural odorants in the mouse olfactory bulb, Nature Neuroscience, 15(4):537–9, 2012.

[51] C. Su, K. Menuz and J. R. Carlson, Olfactory Perception: Receptors, Cells, and Circuits, Cell, 139(1):45–59, 2011.

[52] K. J. Grossman et al, Glomerular activation patterns and the perception of odorant mixtures, European Journal of Neuroscience, Vol. 27, pp. 2676–2685, 2008.

[53] D. A. Wilson and R. M. Sullivan, Cortical Processing of Odor Objects, Neuron, 72(4):506–19, 2011.

[54] K. I. Nagel and R. I. Wilson, Biophysical mechanisms underlying olfactory receptor neuron dynamics, Nature Neuroscience, 14(2):208–16, 2011.

[55] J. Del Castillo and B. Katz, Quantal Components of the End-Plate Potential. J. Physiol, 124, 560–573, 1954.

[56] P.G. Strange, Agonist binding, agonist affinity and agonist efficacy at G protein-coupled receptors. British Journal of Pharmacology, 153(7), 1353–1363, 2009.

[57] M.L. Fletcher, Analytical Processing of Binary Mixture Information by Olfactory Bulb Glomeruli, PLOS One, 6(12), 2011.

[58] R.C. Li, Y. Ben-Chaim, K.W. Yau and C.C. Lin, Cyclic-nucleotide-gated cation current and Ca^2+^-activated Cl^−^ current elicited by odorant in vertebrate olfactory receptor neurons, Proc. Natl. Acad. Sci., 113(40):11078–11087, 2016.

[59] D. Munch, B. Schmeichel, A.F. Silbering and G.C. Galizia. Weaker ligands can dominate an odor blend due to syntopic interactions. Chemical Senses, 38(4), 293–304, 2013.

[60] Wachowiak, M., McGann, J. P., Heyward, P. M., Shao, Z., Puche, A. C., and Shipley, M. T, Inhibition of olfactory receptor neuron input to olfactory bulb glomeruli mediated by suppression of presynaptic calcium influx. Journal of Neurophysiology, 94(4), 2700–2712, 2005.

[61] McGann, J. P., Pirez, N., Gainey, M. A., Muratore, C., Elias, A. S., and Wachowiak, M, Odorant representations are modulated by intra-but not interglomerular presynaptic inhibition of olfactory sensory neurons. Neuron, 48(6), 1039–1053, 2005.

[62] X. Grosmaitre, A. Vassalli, P. Mombaerts, G.M. Shepherd and M. Ma. Odorant responses of olfactory sensory neurons expressing the odorant receptor MOR23: a patch clamp analysis in gene-targeted mice. Proc. Natl. Acad. Sci. USA, 103(6), 1970–1975, 2006.

[63] X. Grosmaitre, X, S.H. Fuss, A.C. Lee, K.A. Adipietro, H. Matsunami, P. Mombaerts and M. Ma. SR1, a mouse odorant receptor with an unusually broad response profile. J Neurosci, 29(46), 14545–52, 2009.

[64] R. C. Challis et al. An olfactory cilia pattern in the mammalian nose ensures high sensitivity to odors. Current Biology 25:2503–12, 2015.

[65] M. Carandini and D.J. Heeger. Normalization as a canonical neural computation. Nature Reviews Neuroscience 13:51–62, 2011.

[66] M. Wachowiak, L.B. Cohen and M.R. Zochowski. Distributed and concentration-invariant spatial representations of odorants by receptor neuron input to the turtle olfactory bulb. J Neurophysiol, 87(2), 1035–1045, 2002.

[67] T.B. Cleland, Johnson, M. Leon and C. Linster. Relational representation in the olfactory system. Proc. Natl. Acad. Sci. USA, 104(6), 1953–1958, 2007.

[68] S.R. Olsen, V. Bhandawat and R.I. Wilson. Divisive normalization in olfactory population codes. Neuron 66, 287–299, 2010.

[69] B. Roland et al. Massive normalization of olfactory bulb output in mice with a monoclonal nose. eLife, 5, 2016.

[70] C. F. Stevens, A statistical property of fly odor responses is conserved across odorants, Proc. Natl. Acad. Sci, 113(24):6737–6742, 2016.

[71] P.T. Wojcik and Y.B. Sirotin. Single scale for odor intensity in rat olfaction. Current Biology, 24(5), 568–573, 2014.

[72] B. Berglund, U. Berglund, T. Engen and G. Ekman, Multidimensional analysis of twenty-one odors, Scandinavian Journal of Psychology, 14(1)131–137, 1973.

[73] D.G. Laing, H, Panhuber, M.E. Willcox and E.A. Pittman. Quality and intensity of binary odor mixtures. Physiol Behav 33, 309–319, 1984.

[74] B. Berglund and M.J. Olsson, Odor-intensity interaction in binary and ternary mixtures. Perception & Psychophysics, 53(5)475–482, 1993.

[75] Kapoor V., Provost A.C., Agarwal P., and Murthy VN, Activation of raphe nuclei triggers rapid and distinct effects on parallel olfactory bulb output channels. Nature Neuroscience, 19:271–282, 2016.

[76] V. Bhandawat, J. Reisert and K. Yau, Signaling by olfactory receptor neurons near threshold, Proc. Natl. Acad. Sci, 107(43):18682–18687, 2010.

[77] R. J. Flannery, D. A. French, S. J. Kleene, Clustering of cyclic-nucleotide-gated channels in olfactory cilia. Biophysical Journal, 91(1):179–188, 2006.

[78] H. Takeuchi, T. Kurahashi, Mechanism of signal amplification in the olfactory sensory cilia, J. Neurosci, 25(48):11084–11091, 2005.

[79] J. Zheng, W. N. Zagotta, Stoichiometry and assembly of olfactory cyclic nucleotide-gated channels. Neuron, 42:411–21, 2004.

[80] I. H. Segel, Enzyme Kinetics: Behavior and Analysis of Rapid Equilibrium and Steady-State Enzyme Systems, Wiley-Interscience, 1993.

[81] J. Reisert, J. Lai, K.W. Lau, J. Bradley, Mechanism of the excitatory Cl- response in mouse olfactory receptor neurons, Neuron, 45(4):553–61, 2005.

[82] T.Y. Chen, H. Takeuchi and T. Kurahashi, Odorant inhibition of the olfactory cyclic nucleotide-gated channel with a native molecular assembly, J. Gen. Physiol., 128(3):365–71, 2006.

